# Automated customization of large-scale spiking network models to neuronal population activity

**DOI:** 10.1101/2023.09.21.558920

**Authors:** Shenghao Wu, Chengcheng Huang, Adam Snyder, Matthew Smith, Brent Doiron, Byron Yu

## Abstract

Understanding brain function is facilitated by constructing computational models that accurately reproduce aspects of brain activity. Networks of spiking neurons capture the underlying biophysics of neuronal circuits, yet the dependence of their activity on model parameters is notoriously complex. As a result, heuristic methods have been used to configure spiking network models, which can lead to an inability to discover activity regimes complex enough to match large-scale neuronal recordings. Here we propose an automatic procedure, Spiking Network Optimization using Population Statistics (SNOPS), to customize spiking network models that reproduce the population-wide covariability of large-scale neuronal recordings. We first confirmed that SNOPS accurately recovers simulated neural activity statistics. Then, we applied SNOPS to recordings in macaque visual and prefrontal cortices and discovered previously unknown limitations of spiking network models. Taken together, SNOPS can guide the development of network models and thereby enable deeper insight into how networks of neurons give rise to brain function.

## Introduction

Computational models facilitate our understanding of brain function by attempting to reproduce specific aspects of the brain’s activity. Single-neuron models, such as the Hodgkin-Huxley model^1^ have provided a mechanistic foundation for the generation of action potentials. Small neural circuit models, such as the stomatogastric ganglion (STG) model of crustaceans^2^, have been used to understand the mechanisms underlying the generation of rhythmic motor patterns. At the systems level, large-scale network models, including rate-based recurrent neural networks^3–5^ and convolutional neural networks^6^, have been highly influential in elucidating how neural circuits perform complex brain computations. Although neurons communicate with each other through temporally complex spike trains, these network models focus on replicating neuronal firing rates without spikes. To establish a link between computational models and biological spiking neurons, large-scale spiking neural networks (SNN) have been proposed. These SNNs aim to produce population spike trains whose time course and/or variability mimic that of neuronal recordings^7–12^. SNNs have become an increasingly important class of large-scale models in computational neuroscience: studying the mechanisms of the biologically realistic circuit of a SNN is a critical step in understanding complex processing in cortical circuits.

A key goal in constructing network models is to customize their parameters to recapitulate some aspect of the recorded neuronal activity. In single-neuron models and small neural circuit models, each model parameter corresponds to a specific biological component and can often be measured experimentally^1, 2^. Larger-scale models, including rate networks and SNNs, are more difficult to customize because of the larger parameter space that comes with a larger number of neurons^13, 14^. Furthermore, comparison between model activity and neuronal recordings is challenging because there is typically not a one-to-one correspondence between each neuron in the model network and each recorded neuron^15^. In particular, the number of neurons within the model network is often far smaller than the number of neurons that comprise the biological network that the model is intended to describe.

Different approaches have been used to circumvent the need for a one-to-one correspondence between model neurons and recorded neurons. One approach is to construct a network model whose output, such as a limb movement^9, 16^ or a decision^17^, reproduces a subject’s behavior given a network input. In such cases, the models are customized by optimizing a cost function representing the difference between the model output and the behavior. Once customized, these network models have shown impressively similar network activity features to activity recorded in the brain, albeit without explicit matching of neuronal activity in the cost function. The cost function can be optimized using methods such as FORCE^13^ or backpropagation^18, 19^ because it has a closed-form expression with respect to the model parameters.

Another approach is to reproduce statistical measures of the neuronal activity. SNNs are often designed to reproduce variability in the activity of individual neurons (e.g., Fano factor of spike counts^20–23^) and covariability between neurons (e.g., pairwise correlation of spike counts^14, 24–26^). SNNs have also been designed to reproduce population-wide covariability in neuronal recordings^27^. The cost function, representing the difference in spiking activity statistics between the model and recordings, has no closed-form expression with the model parameters because it depends on computationally demanding numerical simulations and cannot be directly evaluated. To date, the parameters of these SNNs have been hand-tuned^27^, customized using exhaustive search^14, 28^, or customized using deep learning approaches when the network simulation time is small^29^. This has limited the exploration and understanding of the full range of activity regimes that large-scale SNNs are capable of exhibiting.

Here we propose an automatic framework, Spiking Network Optimization using Population Statistics (SNOPS), for customizing the parameters of a large-scale SNN to reproduce observed spiking activity statistics. SNOPS uses Bayesian optimization^30^ to determine model parameters, a technique widely applied in machine learning for optimizing cost functions without a closed-form expression. We include population-wide activity statistics based on dimensionality reduction to obtain a closer match of the network model to neuronal recordings than using statistics defined only on individual neurons and pairs of neurons^31, 32^. SNOPS provides a guided search of the parameter space and can help accelerate the development of SNNs, especially in settings where the network simulation time is large.

In the following sections, we first introduce the activity statistics used in this work and the SNOPS optimization framework. We next validate SNOPS using activity from a classical balanced network (CBN)^7, 33–35^, the most widely-studied SNN. As a case study, we then apply SNOPS to customize the CBN and its extension, the spatial balanced network (SBN)^11, 27^, to macaque visual area V4 and prefrontal cortex (PFC) recordings. We reveal that SBNs are better suited to reproduce key aspects of neuronal recordings than CBNs. We further identify the performance bottleneck of the CBN by finding the specific combinations of activity statistics that the CBN cannot capture well. Our work provides an automatic framework to close the gap between large-scale spiking network models and large-scale recordings, thereby furthering our understanding of the network-level mechanisms underlying brain function.

## Results

Spiking network models have often been used to generate neuronal activity that resembles that recorded in the brain. Depending on which combination of parameters is chosen, a network can produce spike trains with diverse properties. In a CBN, which consists of recurrently connected excitatory and inhibitory neurons that lack any ordering in their synaptic projections (see Methods), four regimes of population spiking activity have been previously identified based on if the neurons fire with temporal regularity and if neurons are synchronized across the population^34^ (Fig. 1a). This complexity, while being a feature, presents some clear challenges in model customization to recordings. Specifically, it is difficult to search the high-dimensional parameter space to find a set of parameters that produces spike trains with specified properties (e.g., to reproduce some aspects of neuronal population recordings). In this work, we parameterized the CBN using 9 parameters, which govern the connection strength between neurons as well as the timescales of synaptic decay (see Table 1 in Methods). Simulating networks using all possible combinations of parameters can be computationally intractable even with 9 parameters due to the exponential growth in the number of combinations with the number of parameters. Furthermore, it is unknown a priori whether there even exists a combination of parameters (referred to as a *parameter set*) for a given network model that produces spiking activity with the specified properties. Hence there is a clear need of an automated framework to search the parameter space.

**Table 1:** Free parameters for both the CBN and SBN.

| Parameter description | Symbol | Search range |
| --- | --- | --- |
| recurrent inhibitory synaptic decay time constant | $\tau^{id}$ | 1 to 25 ms |
| recurrent excitatory synaptic decay time constant | $\tau^{ed}$ | 1 to 25 ms |
| recurrent I to E connection strength | $J^{ei}$ | -150 to 0 mV |
| recurrent E to I connection strength | $J^{ie}$ | 0 to 150 mV |
| recurrent I to I connection strength | $J^{ii}$ | -150 to 0 mV |
| recurrent E to E connection strength | $J^{ee}$ | 0 to 150 mV |
| feedforward to recurrent E connection strength | $J^{eF}$ | 0 to 150 mV |
| feedforward to recurrent I connection strength | $J^{iF}$ | 0 to 150 mV |

**Figure 1.**
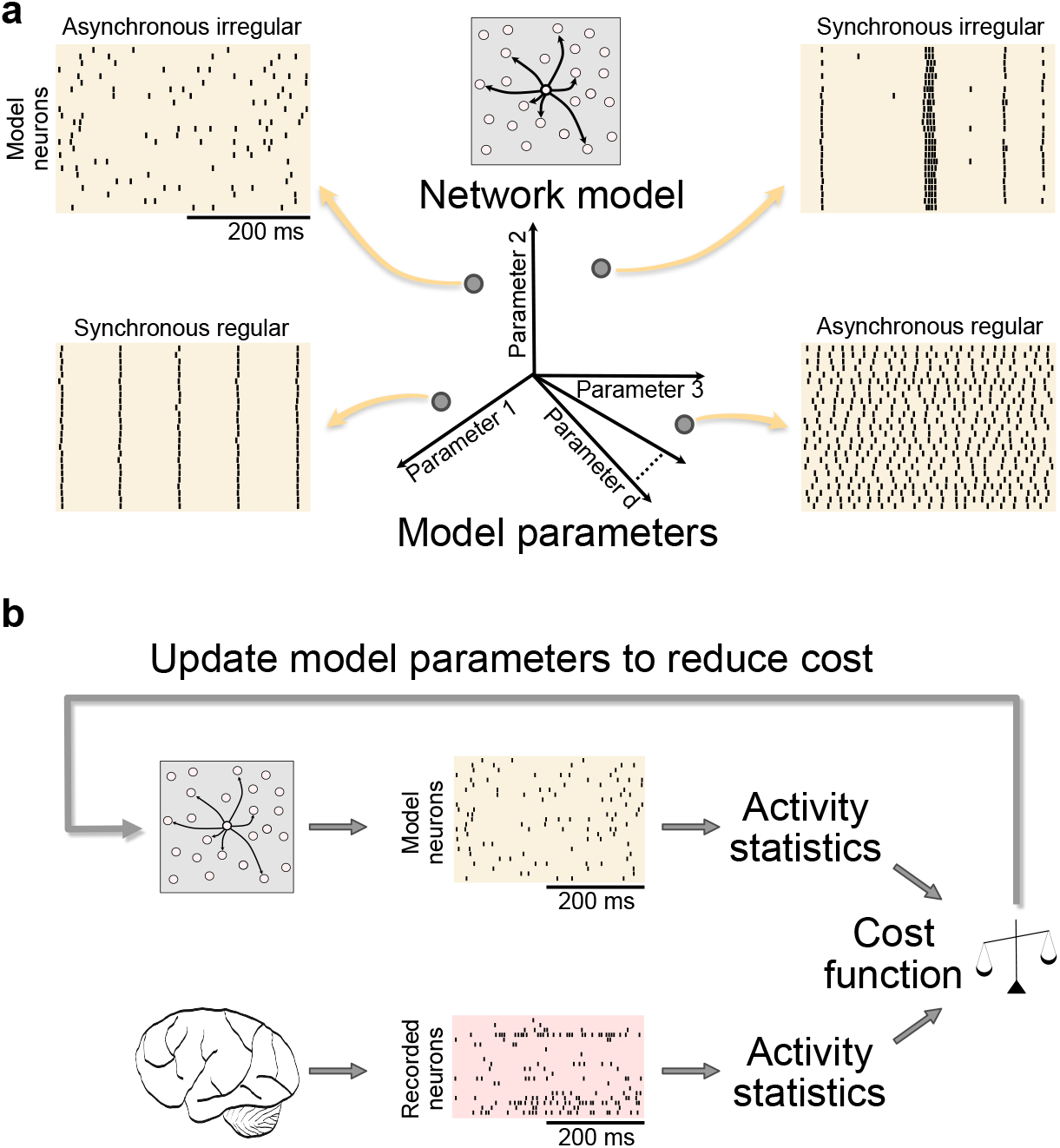
Framework for automated customization of a spiking network model to neuronal recordings. **a**, A SNN has a complicated dependency between its parameters and spiking output. For example, different parameter sets correspond to each of four previously-identified activity regimes of a classical balanced network: asynchronous irregular, synchronous regular, synchronous irregular, and asynchronous regular. In this case, the SNN has a 9-dimensional parameter space. **b**, Our customization framework matches activity statistics of spike trains produced by the network model to those of neuronal recordings. It uses a guided searching algorithm to iteratively update the model parameters. The activity statistics are defined by the user and can include single-neuron, pairwise, and population activity statistics.

We propose a framework (SNOPS) to automatically customize a SNN to neuronal recordings (Fig. 1b). It is based on iteratively updating the model parameters to improve the correspondence between the spike trains generated by the network model and the recorded spike trains (using a *cost function*, defined below). The cost function is based on a set of activity statistics, which are computed for both the generated and recorded spike trains.

### Single-neuron, pairwise, and population activity statistics for comparing models to neuronal activity

We first introduce the activity statistics used to compare the spike trains produced by the network model and the recorded spike trains. For all activity statistics, we begin by counting spikes within pre-defined time bins (Fig. 2a, left). This activity from individual neurons recorded simultaneously can be represented in a population activity space, where each axis represents the activity level of one neuron (Fig. 2a, center). We then compute activity statistics based on individual neurons, pairs of neurons, and populations of neurons (Fig. 2a, right), as described below.

**Figure 2.**
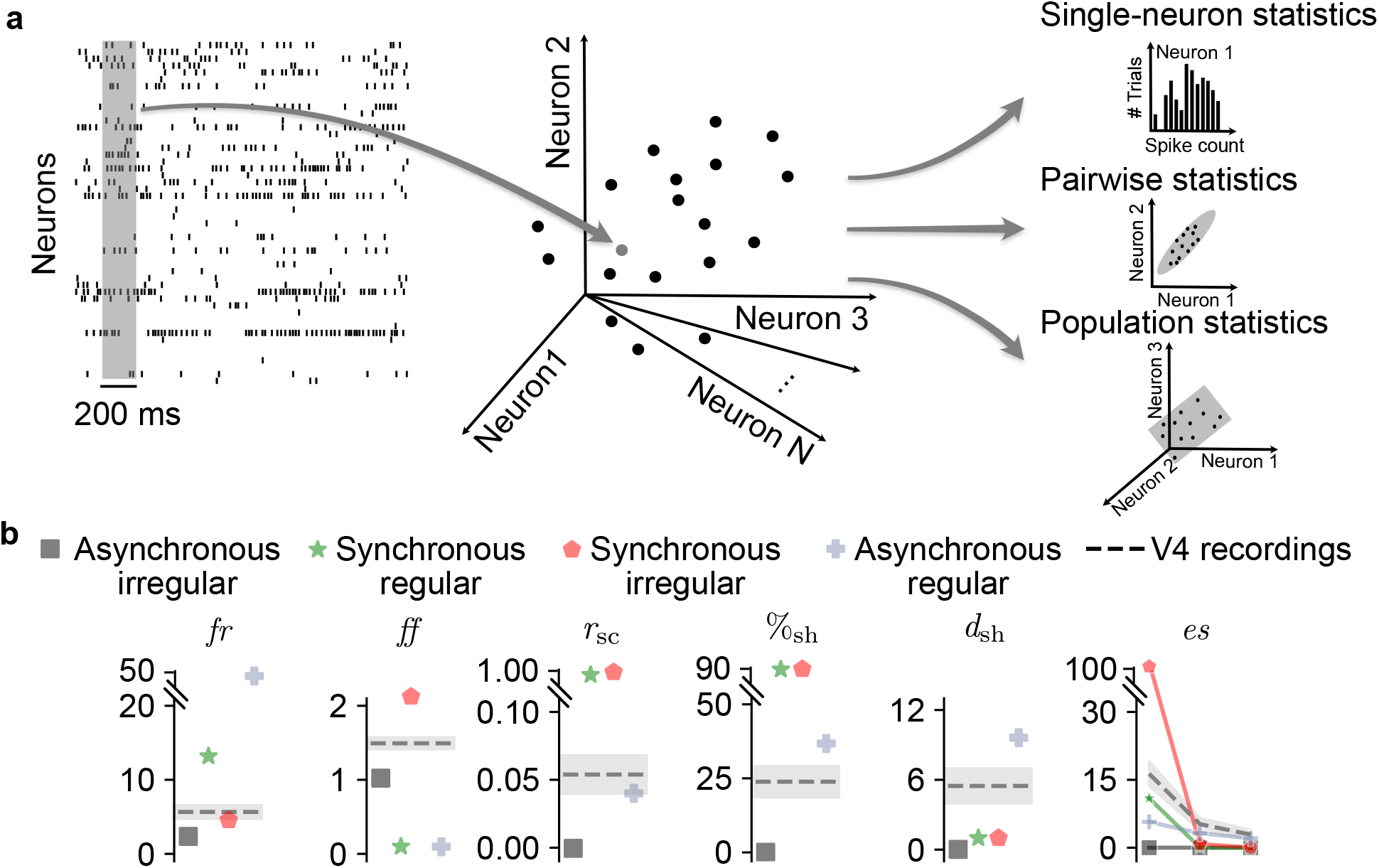
Activity statistics for comparing the activity of a spiking network model to neuronal recordings. **a**, Three types of activity statistics based on single neurons (e.g., firing rate and Fano factor), pairs of neurons (e.g., spike count correlation), and a population of neurons (e.g., percent shared variance, number of dimensions, and eigenspectrum of shared variance). The units of firing rate are spikes per second, those of Fano factor are spike count, and those of the eigenspectrum of shared variance are (spike count)^2^. All other activity statistics are unitless. These activity statistics are all based on spike counts within a 200 ms spike count bin (left panel), which can be represented in a population activity space (center panel). Each dot represents the activity across the neuronal population within a given time window. **b**, Activity statistics based on population recordings in macaque visual area V4 (dashed lines) are challenging to reproduce by the four parameter regimes of a CBN (colored symbols, cf. Fig. 1a, mean across 5 network instantiations of network connectivity graphs and initial membrane potentials corresponding to the same network parameter set). None of the four activity regimes accurately reproduces the activity statistics of the V4 population recordings (dashed lines). The V4 activity statistics are shown as the mean*±*1 SD across 19 recording sessions (see Methods). All activity statistics are based on randomly subsampling 50 neurons from each CBN or V4 dataset.

We considered two single-neuron statistics: mean firing rate (*fr*) and Fano factor (*ff*) (Fig. 2b). The mean firing rate is defined as the average level of activity across all neurons in the population and across all time bins. The Fano factor captures the activity variability of individual neurons across time^36^. For the pairwise statistic, we computed the spike count correlation between pairs of neurons (*r*_sc_, Fig. 2b), which is widely used to measure the correlated variability among neurons^37^. Both single-neuron and pairwise statistics have been widely used to customize network models to neuronal recordings^9, 13, 17, 21, 27, 38–40^.

There can also be structure in the population-wide variability that is not apparent when considering only single-neuron and pairwise statistics^32^. Previous studies have used population-wide activity statistics to compare network models to recorded activity^27, 31, 41, 42^. Thus we also considered population activity statistics based on dimensionality reduction^43^. Specifically, we used factor analysis (FA), which is the most basic dimensionality reduction method that separates the variance that is shared among neurons from the variance that is independent to each neuron. We computed the following three population activity statistics based on FA (Fig. 2b; see Methods and Supplementary Fig. 1): (1) The percent shared variance (%_sh_) is the fraction of a neuron’s activity variance that is shared with one or more of the other neurons in the recorded population. This value is first computed per neuron, then averaged across neurons. A high %_sh_ indicates that the population of neurons strongly covary, whereas zero %_sh_ indicates that neurons are independent of each other. While %_sh_ is related to *r*_sc_, it is not identical and captures a different aspect of population activity^32^. (2) We measured the dimensionality as the number of dimensions needed to explain the shared variance among neurons (*d*_sh_). If the neurons all simply increase and decrease their activity together, *d*_sh_ would equal one. If the neurons covary in more complex ways, *d*_sh_ would be greater than one. (3) The eigenspectrum (*es*) of the shared covariance matrix measures the relative dominance of the dimensions identified above. It may be that the first dimension explains far more shared variance than the other dimensions (in which case the eigenspectrum would have a sharp dropoff) or that all dimensions explain a similar amount of shared variance (in which case the eigenspectrum would be flat).

### Manual customization of SNNs to neuronal activity can be challenging and labor intensive

The art of manually customizing a SNN to neuronal recordings is fraught: it can be difficult to intuit the resulting activity of the network from changes to its parameters. To get around this difficulty, one might ask whether any of the four CBN activity regimes shown in Fig. 1a capture the key aspects of neuronal recordings (Fig. 2a, left). We thus computed the single-neuron, pairwise, and population activity statistics of the spike trains shown in Fig. 1a for each of the four activity regimes (Fig. 2b, colored shapes). We compared them to the activity statistics computed from the population activity recorded in macaque visual area V4 (Fig. 2b, dashed lines). We found that none of the four previously-identified CBN activity regimes recapitulated all of the activity statistics of the V4 neurons. Thus, we need an automatic method to search the parameter space to determine if there exists a parameter set whose activity better resembles neuronal recordings.

### Customizing SNNs using Bayesian optimization

The central contribution of this work is an automatic framework, Spiking Network Optimization using Population Statistics (SNOPS, Fig. 3), to address this need. SNOPS iteratively updates the parameters of the SNN so that the activity statistics of the model-generated activity (Fig. 3a) better match those of the recorded neuronal activity (Fig. 3b). To quantify how well matched are the two sets of activity statistics, we define a cost function (Fig. 3c) as a linear combination of the squared difference between the two sets of activity statistics (see Methods).

**Figure 3.**
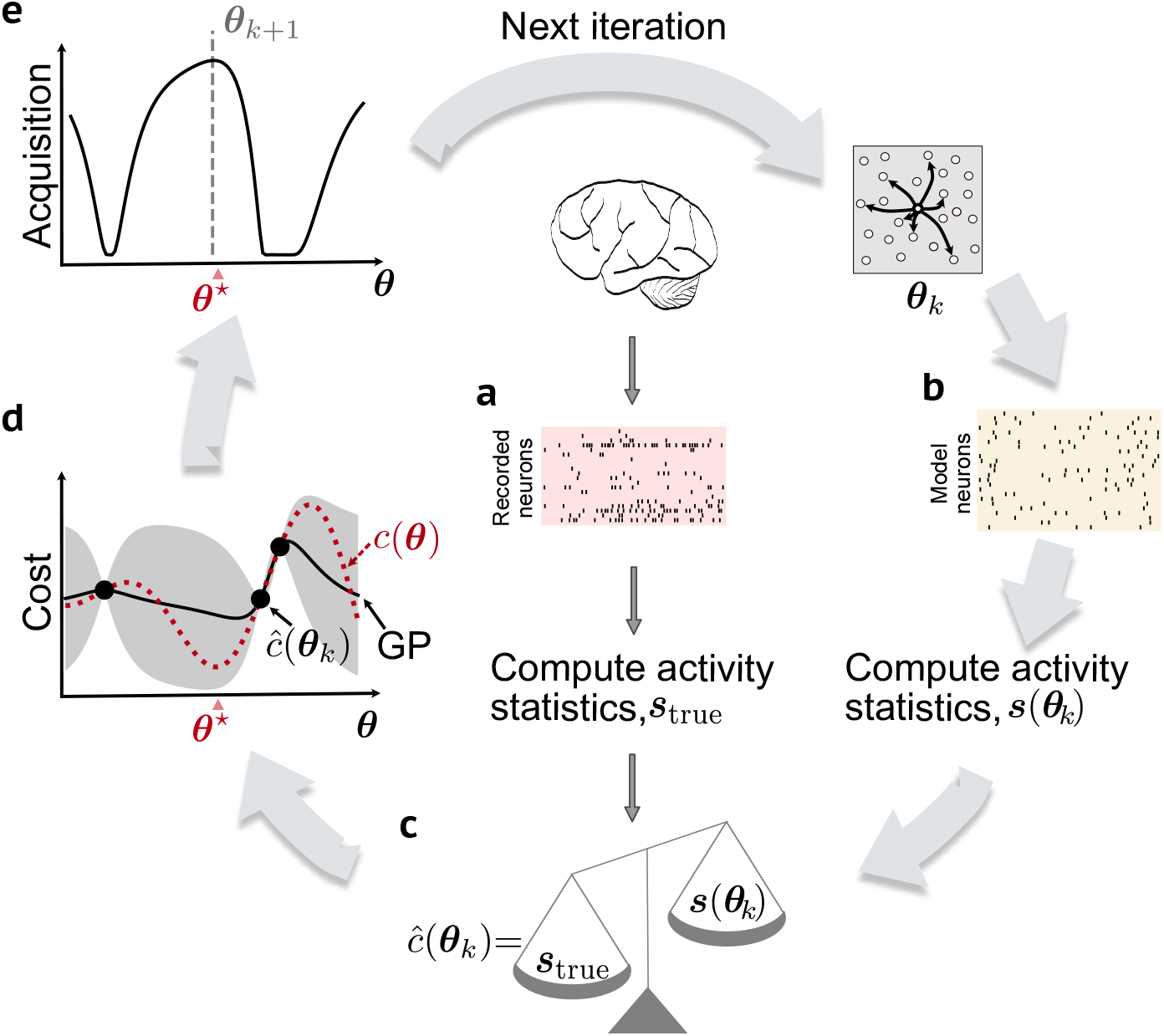
Customizing a spiking network model using Bayesian optimization with Gaussian processes. The Bayesian optimization algorithm attempts to find a parameter set ***θ*** for a spiking network model such that its activity statistics match those of neuronal recordings. **a**, Spike trains are recorded from the brain and their activity statistics, ***s***_true_, are computed. This step is performed only once, since the same recorded activity is used for comparison on all iterations. **b**, On the *k*-th iteration, spike trains are generated from the network model using parameter set ***θ***_*k*_, proposed by the previous iteration. **c**, The activity statistics of the spike trains generated from the network model, ***s***(***θ***_*k*_), are computed. The cost for ***θ***_*k*_ depends on how far each of those activity statistics is from the corresponding activity statistics of the neuronal recordings, ***s***_true_. **d**, A Gaussian process (GP) (solid line) is used to approximate the true, unknown cost function, *c*(***θ***) (red dashed line). We seek to find the minimum of this true, unknown cost function (denoted by ***θ***^*\**^). Each iteration of the Bayesian optimization provides one evaluation of the cost at a particular setting of the model parameters (black dots). The cost at the current iteration is labeled (***θ***_*k*_), and the other black dots represent the costs evaluated during previous iterations. The GP provides an uncertainty of our estimate of the cost function (gray shading). For illustrative purposes, we show here a single model parameter being optimized, whereas our algorithm typically optimizes multiple model parameters simultaneously. **e**, An acquisition function is defined based on the GP in **d** to determine the next parameter set, ***θ***_*k*+1_, to evaluate. The acquisition function implements an exploration-exploitation trade-off, where areas of low predicted cost and high uncertainty are desirable.

Assessing how adjusting any model parameter influences the cost requires generating spiking activity from the network model. Therefore, the cost function cannot be expressed in a closed form with respect to the model parameters and cannot be optimized using gradient methods. Meanwhile, exhaustive search methods, such as random search, may yield excessive running time because it is computationally demanding for the network model to generate spikes. Instead, we need a guided way of searching the parameter space. Bayesian optimization is a natural choice to optimize a cost function whose evaluation depends on a time-consuming simulation^44, 45^. It automatically proposes the next model parameter set to evaluate based on the cost of the previously evaluated parameter sets. The key idea is that more similar model parameter sets should correspond to more similar costs. This relationship is described by a Gaussian process (GP)^46^. The algorithm uses the GP to propose model parameters whose predicted cost is low (i.e., parameter sets that may be better than those already considered) and whose uncertainty about the predicted cost is high (to sample from unvisited areas of the parameter space). This defines an exploitation-exploration trade-off.

The GP (Fig. 3d, solid line) approximates the cost function *c*(***θ***) (Fig. 3d, red dashed line), which is a priori unknown, using all evaluations of the cost function from previous iterations and the current iteration (Fig. 3d, dots). Bayesian optimization will then construct an acquisition function (Fig. 3e) based on the GP-predicted cost and its uncertainty at each setting of the model parameters ***θ*** (see Methods). The parameters ***θ***^*\**^ that maximize the acquisition function are selected for the next iteration, and the entire process restarts (i.e., the new parameter set ***θ***^*\**^ is used to simulate spike trains from the SNN, whose activity statistics are then computed, etc.). With more iterations, Bayesian optimization will likely sample parameter sets with lower cost, until a stopping criterion has been reached (see example in Supplementary Fig. 2). To further accelerate the customization procedure, we introduced two computational innovations in SNOPS: (1) running a short simulation to assess whether a parameter set is likely to yield valid spike trains (feasibility constraint, see Methods), and (2) dynamically increasing the number of simulations to reduce the variance of the estimated cost (intensification, see Methods).

### SNOPS accurately recovers activity statistics in simulation

To validate SNOPS, we first generated activity from a CBN and computed its activity statistics (Fig. 4a). These served as the target activity statistics in the customization procedure, in place of activity statistics computed from neuronal recordings. We then used SNOPS to customize a separate CBN to these activity statistics. In this case, there is no model mismatch. Thus, there exists a CBN parameter set that reproduces the target activity statistics exactly.

**Figure 4.**
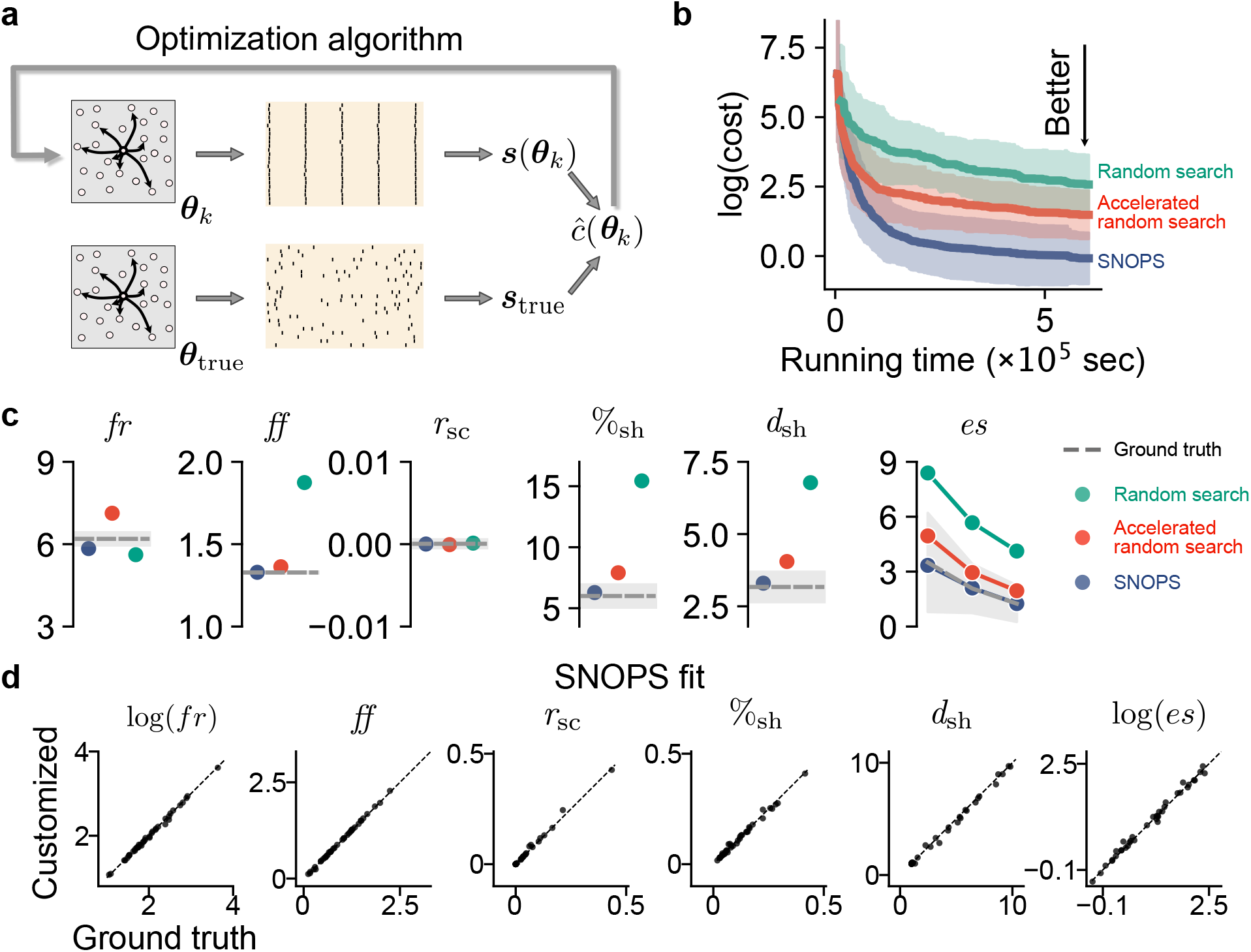
SNOPS accurately customizes a classical balanced network model to simulated spike trains. **a**, A CBN was used to generate spike trains with randomly chosen parameter sets ***θ***_true_ (see Methods). SNOPS (or other optimization algorithms) was then used to customize the parameters, ***θ***_*k*_, of a separate CBN to match the “ground truth” activity statistics, ***s***_true_, of the generated spike trains. **b**, For a given amount of computer running time (see Methods), SNOPS (blue) finds parameters with lower cost than accelerated random search (red) and random search (green). Vertical axis represents the lowest log(*cost*) up to the given running time and hence decreases monotonically. Solid lines and shading represent the mean*±* 1 SD across 40 customization runs. **c**, For a representative customization run, SNOPS (blue) identified model parameters whose activity statistics were closer to the ground truth (dashed lines) than accelerated random search (red) and random search (green). Error bars on the ground truth represent one SD across 5 network instantiations corresponding to the same ground truth parameter set. Circles represent the mean across 5 network instantiations corresponding to the network parameter set identified by each optimization algorithm. **d**, Across all 40 customization runs, SNOPS accurately reproduced the ground truth activity statistics (all points lie near the diagonal). Each dot represents the results from one SNOPS customization run to a randomly generated ground truth dataset. For visual clarity, only the first (i.e., most dominant) mode of *es* is plotted in the rightmost panel.

For comparison, we repeated the customization task with two other optimization algorithms applicable to large-scale SNNs which do not have a closed-form cost function: random search and its accelerated variant. Random search proceeds by sampling parameter sets from the search region uniformly at random. This method is similar to the exhaustive search approach in previous literature^14^ and provides a benchmark for performance comparisons. The accelerated random search incorporates two computational innovations that we introduced in SNOPS (feasibility constraint and intensification, see Methods). Therefore, when going from random search to its accelerated variant, the only difference is incorporating the two innovations in SNOPS. Going from accelerated random search to SNOPS, the only difference is replacing random search with Bayesian optimization. This arrangement enables us to systematically qualify the benefits of the two key features of SNOPS: Bayesian optimization and the two innovations.

SNOPS (Fig. 4b and Supplementary Fig. 3, blue) outperformed accelerated random search (Fig. 4b and Supplementary Fig. 3, red), indicating that Bayesian optimization achieves a lower cost than random search after the same amount of computer running time. Accelerated random search outperformed random search (Fig. 4b and Supplementary Fig. 3, green), indicating that the two innovations of SNOPS are beneficial. Furthermore, all three methods yielded a CBN whose activity statistics better match the target activity statistics with increasing running time, as expected (Fig. 4b and Supplementary Fig. 3). Another related method, Sequential Neural Posterior Estimator (SNPE)^29^, returns a distribution of parameter sets and requires generating a large number of SNN simulations upfront. We compare SNOPS to SNPE in Supplementary Fig. 4 (also see Discussion).

The cost (vertical axis in Fig. 4b) is a summary of how accurately the activity statistics of the customized CBN match the target activity statistics. To further understand the difference in performance between these methods, we then compared the individual activity statistics returned by each method to their target values. Consistent with Fig. 4b, SNOPS was better able to match the target activity statistics than the other methods (Fig. 4c). Across all 40 customization runs, SNOPS successfully identified CBNs whose activity statistics closely matched the target activity statistics (Fig. 4d, all the dots are located near the diagonal). In sum, SNOPS customizes a spiking network model to neuronal activity more quickly and accurately than the other methods.

### Case study: customizing SNNs to V4 and PFC population recordings using SNOPS

We next present a case study of using SNOPS to customize SNNs to neuronal population recordings in macaque monkeys (from Utah arrays implanted in visual cortical area V4 and in prefrontal cortex, or PFC).

In Fig. 2b, we demonstrated that none of the four CBN activity regimes from Fig. 1a recapitulated the V4 datasets (mean cost*±*SD, asynchronous irregular: 13.56*±*0.12, synchronous regular: 1823.67*±*190.06, synchronous irregular: 1489.71*±*124.25, asynchronous regular: 361.44*±*9.30). Here we used SNOPS to automatically customize a CBN to the same V4 datasets and obtained a substantially lower cost (2.71*±*0.24, *p <* 1 *×* 10^*−*5^ for each of the four comparisons, one-sided t-test). In other words, SNOPS reproduced the activity statistics more accurately than any of the four previously-identified activity regimes (Fig. 5a, compare to Fig. 2b).

**Figure 5.**
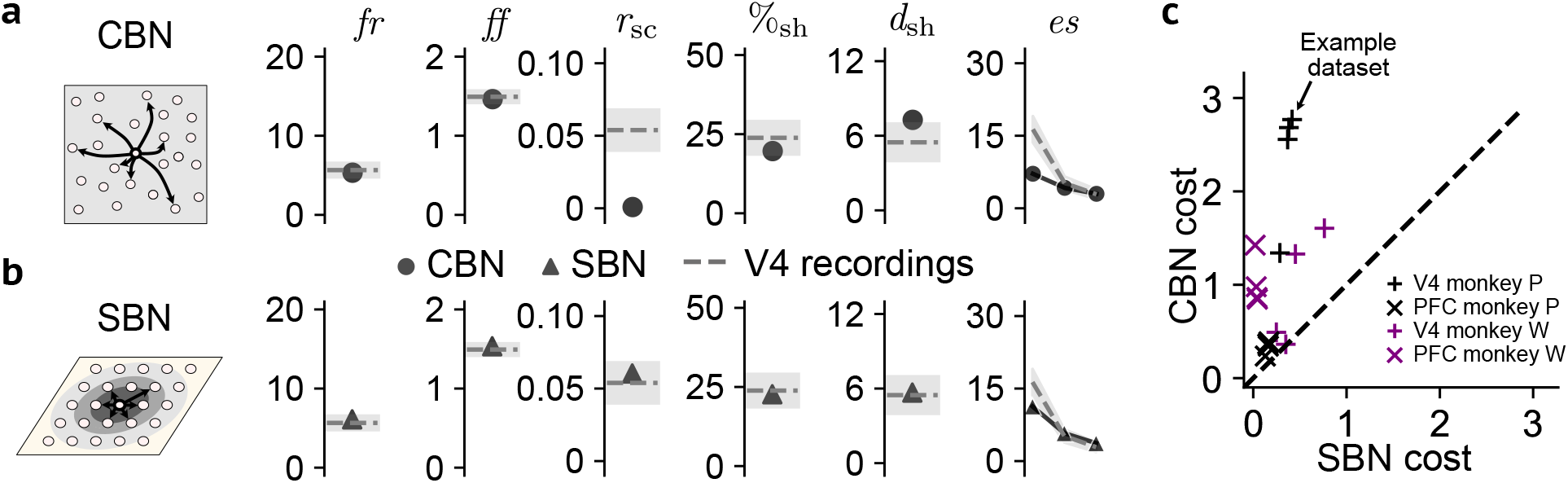
The spatial balanced network (SBN) more accurately reproduces activity statistics of macaque V4 and PFC datasets than the classical balanced network (CBN). **a**, Left: Stylized representation of the CBN. Right: Activity statistics of the CBN (circles, mean across 5 network instantiations corresponding to the same identified parameter set) after being customized using SNOPS to the same V4 dataset as in Fig. 2b. Dashed line and shading represent the mean*±*1 SD across 19 sessions. **b**, Left: Stylized representation of the SBN. The SBN is different from the CBN in that the connection probability depends on the distance between neurons. Right: Activity statistics of the SBN (triangles, mean across 5 network instantiations corresponding to the same identified parameter set) after being customized using SNOPS to the same V4 datasets as in **a. c**, The SBN more accurately reproduced activity statistics than the CBN across 16 datasets, comprising four task conditions with recordings in two brain areas (V4 and PFC) in each of the two monkeys. Arrow indicates the example V4 dataset shown in **a** and **b**.

Despite this improvement, there were activity statistics that were not accurately reproduced. Specifically, the *r*_sc_ for the customized CBN was substantially smaller than that of the V4 datasets (mean SD, 0.00085*±* 0.0011 versus 0.054*±*0.015, *p <* 1*×* 10^*−*4^, one-sided t-test) (Fig. 5a). To verify the reliability of this disagreement, we reran SNOPS with different initializations and obtained the same disagreement in *r*_sc_ (Supplementary Fig. 5). This indicates that this result was not due to the underperformance of SNOPS.

We did not observe such a disagreement in *r*_sc_ when we customized the CBN to the simulated activity generated by CBNs across a wide range of model parameters (Fig. 4d, third panel). This led us to hypothesize that the CBN model framework is not flexible enough to capture the full complexity of the V4 datasets, as measured by the six activity statistics. We thus explored a more powerful SNN model with the goal of more accurately capturing the properties of spiking activity in the V4 datasets.

The spatial balanced network (SBN), an extension of the CBN, has been recently proposed as a SNN framework capable of producing a wider range of population statistics^11, 27^. Neurons in a SBN are organized over a two-dimensional spatial lattice and have connection probabilities that depend on the distance between neuron pairs. This introduces two additional model parameters, for a total of 11 parameters (see Methods). This is in contrast to a CBN model which lacks spatial connectivity and has connection probabilities that are the same for all neuron pairs. SBNs have been heuristically shown to produce activity that resembles the V4 population activity^27^. However, this claim has not been quantitatively verified. We next used SNOPS to systematically explore the capacity for SBNs to capture a wider range of population activity.

We first verified that SNOPS can accurately customize a SBN to simulated activity (Supplementary Figs. 5b and 6), mirroring our results with the CBN (Supplementary Fig. 5a and Fig. 4d). We then customized a SBN to V4 population activity and found that a SBN is able to more accurately reproduce the activity statistics of the V4 population recordings than a CBN (Fig. 5b, mean cost*±* SD, 0.26*±*0.10 versus 2.71*±*0.24, *p <* 1*×*10^*−*5^, one-sided t-test). To test if the benefit of the SBN over CBN is data specific, we customized both models to 16 “datasets”, comprising four task conditions with recordings in two brain areas (V4 and PFC) in each of the two monkeys (see Methods). Across these datasets, the SBN consistently outperformed the CBN in reproducing the activity statistics of the neuronal recordings (Fig. 5c).

### Revealing limits of network model flexibility using trade-offs in activity statistics

To understand why the SBN outperforms the CBN, we customized each SNN to each activity statistic individually rather than all six activity statistics together. We found that the CBN was able to accurately reproduce each activity statistic individually, including *r*_sc_ (Fig. 6a). This suggests that the reason why the CBN is unable to reproduce all six activity statistics simultaneously is due to trade-offs between different statistics: adjusting the model parameters to better reproduce one statistic can affect how accurately another statistic is reproduced.

**Figure 6.**
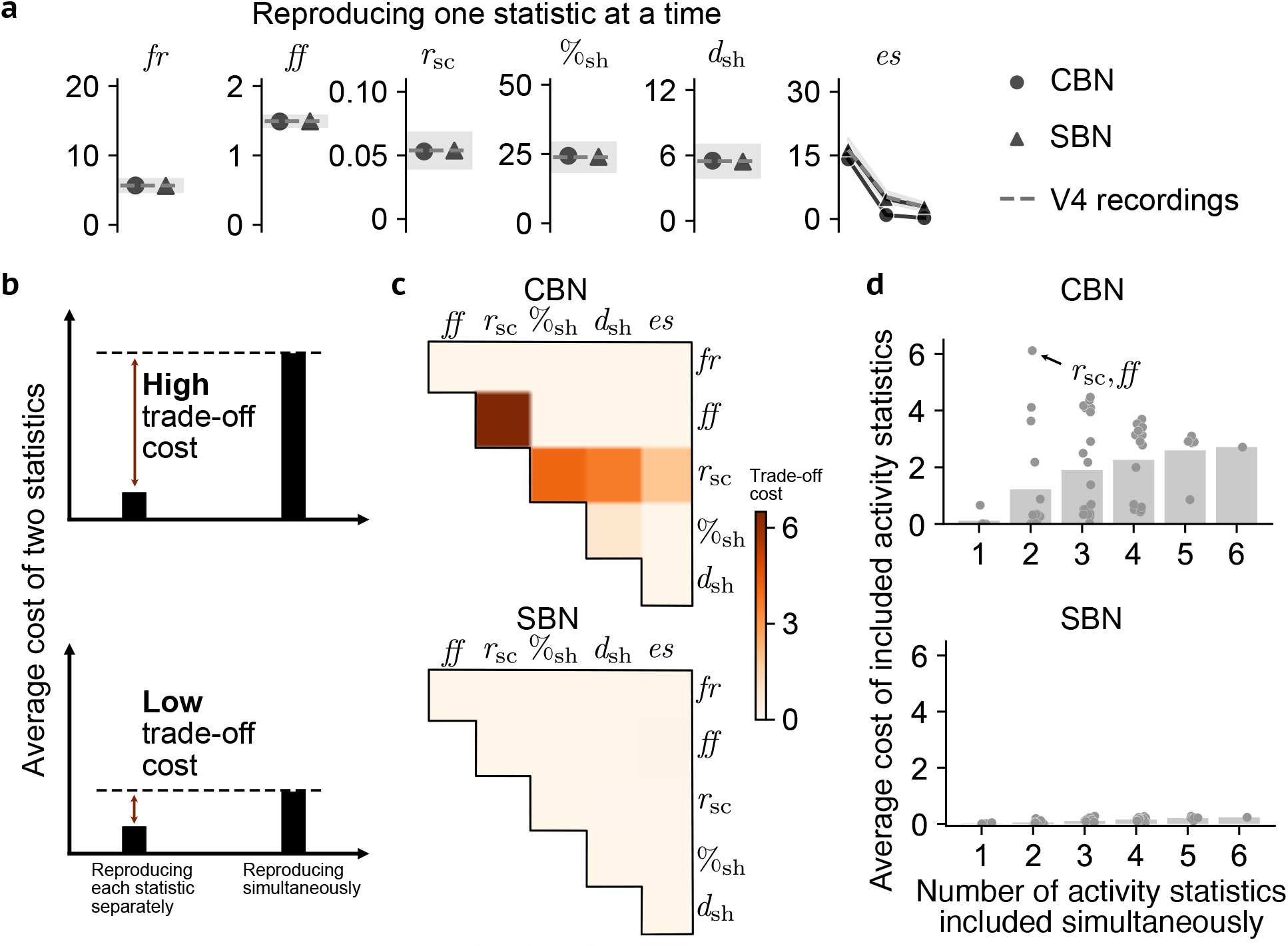
Trade-off cost reveals the inflexibility of CBN relative to SBN. **a**, Activity statistics of the CBN (circles, mean across 5 network instantiations corresponding to the same identified parameter set) and SBN (triangles, mean across 5 network instantiations corresponding to the same identified parameter set) after being customized using SNOPS to one V4 activity statistic (dashed line) at a time. Same V4 dataset as Fig. 2b. **b**, A high trade-off cost represents the case where customizing the network to reproduce two activity statistics simultaneously yields a higher average cost of the two statistics than customizing each statistic individually (upper panel). By contrast, a low trade-off cost represents the case where the cost of customizing two activity statistics simultaneously yields a similar cost to customizing each statistic individually (lower panel). **c**, Trade-off costs between pairs of statistics for the CBN (upper panel) and SBN (lower panel) on the same V4 dataset as Fig. 2b. **d**, Customizing the CBN and SBN to different numbers of activity statistics included in the cost function simultaneously, on the same V4 dataset as Fig. 2b. Each dot represents one particular subset of activity statistics (e.g., highlighted dot indicates the average cost of *r*_sc_ and *ff* when including only those two activity statistics in the cost function). The cost of each dot was computed over 5 network instantiations corresponding to the same identified parameter set of that dot. Each bar indicates the average cost across all subsets of the corresponding number of activity statistics.

We thus defined a *trade-off cost*, which measures whether more accurately reproducing one activity statistic leads to less accurately reproducing another activity statistic. For example, a model might be able to accurately reproduce the %_sh_, but at the expense of making *r*_sc_ too low. In this case, there is a non-zero trade-off cost, indicated by a combined cost of customizing the two statistics simultaneously that is greater than customizing them individually (Fig. 6b). Note that the trade-off cost is distinct from the overall cost, in that an accurate model with a low overall cost might still have a non-zero trade-off cost for particular pairs of statistics.

We used the trade-off cost to understand why the SBN can more accurately reproduce activity statistics of neuronal recordings than the CBN (cf. Fig. 5). We found that the CBN suffers from a trade-off cost between *r*_sc_ and *ff*, as well as between *r*_sc_ and the population statistics (Fig. 6c, upper panel). By contrast, the SBN has a small trade-off cost for all pairs of statistics (Fig. 6c, lower panel). This is due to the flexibility afforded to the SBN by the extra parameters that control the spatial scales of connection probabilities that the CBN lacks (see Methods, the CBN can be a special case of the SBN). A consequence of this flexibility is that the SBN needs to be appropriately constrained during the customization process. For example, if we customize a SBN using only single-neuron and pairwise statistics, the population statistics of the SBN are not accurately reproduced (Supplementary Fig. 7). This demonstrates the value of including population statistics in the customization process, especially for more flexible models like the SBN.

Trade-offs can also occur among more than two statistics. To investigate this, we systematically increased the number of statistics included in the cost function. The average cost of the customized statistics increases as more statistics are included (Fig. 6d). This illustrates how customizing a model to simultaneously reproduce more statistics imposes more constraints on the model customization process. For the CBN, there is already a marked increase in cost when going from one statistic to two statistics included in the cost function (Fig. 6d, upper panel). In particular, there is a high cost of customizing *r*_sc_ and *ff* simultaneously (highlighted dot), consistent with Fig. 6c (upper panel). By contrast, the average cost for the SBN remains low for even up to all six simultaneously customized statistics (Fig. 6d, lower panel). Hence the pairwise trade-off cost we show in Fig. 6c is sufficient for the comparison of model flexibility. In the multi-dimensional space of activity statistics, these trade-offs form a boundary of combinations of statistics that each model is able to reproduce (Supplementary Fig. 8): the tighter boundary of the CBN than the SBN confirms its higher trade-offs as shown in Fig. 6c.

## Discussion

A fundamental goal in computational neuroscience is to develop large-scale SNNs that mimic neuronal activity recorded from the brain. This is a challenging task because the network activity depends on model parameters in a complicated way and generating activity from a SNN is computationally intensive. Our framework for Spiking Network Optimization using Population Statistics (SNOPS) uses Bayesian optimization to guide the parameter search for customizing large-scale SNNs to neuronal activity. We applied SNOPS to customize two SNN models (CBN and SBN) to population recordings from macaque visual and prefrontal cortices. SNOPS revealed that SBNs are more capable of simultaneously reproducing the empirically-observed single-neuron, pairwise, and population activity statistics than CBNs. SNOPS also discovered key limitations of the CBN in reproducing these activity statistics. Overall, SNOPS can guide the development of network models, thereby enabling deeper insights into the circuit mechanisms that underlie brain function.

We emphasize the benefit of incorporating population statistics in our framework to compare the activity of network models and neuronal recordings. Indeed, population statistics have played a key role in characterizing neuronal activity, thereby shedding new light on our understanding of brain function. For instance, population statistics facilitate our understanding of how PFC populations encode color and motion information^17^ as well as working memory^47^, how populations of visual cortical neurons change their activity with visuospatial attention^48, 49^, rhow motor cortical activity changes during learning^50^, and how the same motor cortical neurons can be involved in both movement preparation and execution^51^. Including population statistics in the cost function explicitly encourages a model to reproduce population statistics, which might otherwise not be reproduced if the cost function had only included single-neuron or pairwise statistics (cf. Supplementary Fig. 7).

There are several key considerations when selecting which method to use when customizing a network model: properties of the cost function, the simulation time of the model, and the number of customized models desired for the scientific goal. The first consideration is whether the cost function has a closed-form expression with respect to model parameters. If so, evaluating the cost can be fast, and one can utilize algorithms such as FORCE^13^ to customize the network model. If the cost function is also differentiable with respect to the model parameters (i.e., it has a closed-form gradient), one can customize a network model using methods that utilize the gradient, such as backpropagation^19^ and emergent property inference (EPI)^52^. These approaches are computationally fast and scalable to a large number of parameters. By contrast, the cost function of large-scale SNNs typically has no closed-form expression with respect to the model parameters and hence falls outside the scope of the aforementioned methods. In such cases, three types of algorithms can be used. Evolutionary algorithms, which are biologically inspired, have been applied to customize Hodgkin-Huxley and stomatogastric ganglion models^53, 54^. Sequential Neural Posterior Estimation (SNPE)^29^, a method based on deep generative models, has also been applied to customize these models to find a distribution of the parameter sets whose activity statistics mimic those of the recordings. Finally, Bayesian optimization, as we propose here in SNOPS, can be used to customize large-scale SNNs.

The second consideration is the simulation time of the network model. SNPE requires generating a large number of simulations to train the deep generative model. This is feasible for models with short simulation time, such as Hodgkin-Huxley and stomatogastric ganglion models. Once trained, SNPE can be used to customize the network model repeatedly to a large number of datasets without the need to run additional simulations. This scalability is due to the relationship between model parameters and activity statistics that SNPE learns during training. Large-scale SNNs, however, have a long simulation time due to the large number of neurons in the network, which poses a challenge for simulation-intensive methods such as SNPE. In such settings, optimization-based methods, such as Bayesian optimization, are preferred as they minimize the cost function iteratively without needing to pre-generate a large number of simulations.

The third consideration is the number of customized models desired for the scientific goal. Most studies to date customize a single model to the neuronal recordings (e.g., refs^6, 7, 9, 11, 14, 27^). In this case, SNOPS is preferred because it finds a single customized model more quickly and likely better reproduces the recorded activity than SNPE (cf. Supplementary Fig. 4). SNPE can also be used in this case by selecting the mode of the posterior distribution. However, the running time would be substantially greater for SNPE because it aims to capture the entire distribution of the parameters, which requires substantially more network simulations. Interestingly, there may exist different combinations of parameters that lead to the same network activity^55^. One may seek to interrogate how different parameters compensate each other to produce the same activity^29, 52^. Although SNOPS can be used in this scenario, it requires repeated customization runs with different starting points to obtain multiple solutions, which is computationally demanding. In this case, deep generative models, such as EPI and SNPE, are preferred because they return a parameter distribution characterizing the many parameter sets that lead to the same recorded neuronal activity. However, doing so requires either differentiability (EPI) or the ability to generate a large number of simulations quickly (SNPE).

A major application of SNOPS is to facilitate the development of more flexible models and thereby further our scientific understanding of brain function. This is achieved in the following two ways. First, if certain activity statistics are not accurately reproduced during manual customization, it is unclear whether one needs to continue to manually tune the model parameters in hopes of reproducing all activity statistics, or to consider a new class of models (e.g., by introducing spatial connectivity). SNOPS performs a guided search of the high-dimensional parameter space, thereby providing greater confidence about when a new class of models needs to be considered. Second, the automatic optimization algorithm in SNOPS enables repeated customization of a model with different subsets of activity statistics, facilitating a more complete understanding of a model’s limitations. For example, customization of a CBN to neuronal recordings might suggest that the CBN is incapable of reproducing experimentally-observed *r*_sc_ values (cf. Fig. 5a). In this case, one could be misled to invest time in modifying the network model specifically so that it can reproduce the experimentally-observed *r*_sc_. Using SNOPS, we found that customizing *r*_sc_, in itself, was not problematic. Instead, it was the trade-off between *r*_sc_ and other activity statistics that limited the CBN (cf. Fig. 6c). Such insight will not only profoundly influence plans for making the model more flexible, but also shed light on how different network architectures (e.g., CBN versus SBN) lead to different model flexibility.

Our approach is modular and each component of SNOPS can be tailored towards the scientific goals of the user. First, we can task SNOPS with replicating features of neuronal activity other than those used in this work by incorporating the appropriate activity statistics in the cost function. For example, incorporating timescales of the population activity, such as autocorrelation^56, 57^, may enable a deeper understanding of how circuit structure gives rise to the different timescales of spiking activity across the cortex. We did not include activity timescales in this study because neither SBN nor CBN is designed to capture that aspect of neuronal activity. Second, we can modify the optimization algorithm to speed up the customization process. For example, fitting and predicting with Gaussian processes to approximate the cost function is computationally demanding when the number of parameters is large^46^. One possible solution is to replace Gaussian processes with a more scalable model such as Bayesian neural networks^58^ or neural processes^59^ to reduce the computational time.

Advancements in neuronal recording technologies are enabling measurements of brain activity at unprecedented scale^60–62^. Large-scale models and large-scale neuronal recordings are closely related: large-scale models provide a systematic and mechanistic understanding of large-scale neuronal recordings; whereas large-scale neuronal recordings can further expose limitations of large-scale models. Our automatic algorithm, SNOPS, can be used to accelerate this cycle and facilitate the synergy between model-based (mathematical) approaches and empirical measurements of brain activity to further our understanding of the brain.

## Methods

### Components for customizing a network model

Here we list the components one needs for customizing a network model to neuronal population activity. Each of these components is described in detail in the sections below.

- Neuronal recordings: neuronal activity recorded from a population of neurons (or generated from a network model). In this study, the neuronal activity is in the form of spike trains either recorded experimentally or generated by a spiking network model.
- Network model with unknown parameters: a mathematical model to be customized to the neuronal recordings. In this study, we use a classical balanced network (CBN)^7^ and a spatial balanced network (SBN)^27^.
- Activity statistics: types of activity statistics that are used to measure how similar the activity produced by the network model is to the neuronal recordings. The user can define their own activity statistics depending on their needs. In this study, we use mean firing rate (*fr*), Fano factor (*ff*), spike count correlation (*r*_sc_), percent shared variance (%_sh_), dimensionality (*d*_sh_), and the eigenspectrum of the shared variance (*es*).
- Cost function: a function that takes as input the activity statistics of the network model and those of the neuronal recordings, and outputs a scalar that summarizes how different are the two sets of activity statistics. In this study, we use a weighted sum of squared differences.
- Optimization algorithm: an algorithm for adjusting the parameters of the network model so that its activity resembles the neuronal recordings (i.e., to minimize the cost). In SNOPS, we use Bayesian optimization. Users can also incorporate other optimization algorithms, such as random search and evolutionary algorithms.

### Spiking network models

#### Model details

Since both classical balanced networks (CBNs) and spatial balanced networks (SBNs) are composed of the same single neuron model, we will present both network models together. Each network has one feedforward layer and one recurrent layer. The feedforward layer contains *N*_*F*_ = 625 excitatory neurons emitting spikes according to independent Poisson processes with a uniform rate of 10 spikes per second. There are *N*_*e*_ = 2500 excitatory neurons and *N*_*i*_ = 625 inhibitory neurons in the recurrent layer. Note that this number is smaller than our past work^27^ (where *N*_*F*_ = 2500, *N*_*e*_ = 40, 000, *N*_*i*_ = 10, 000); this was done to reduce the simulation time while maintaining similarly rich network activity. The membrane potential of a neuron *j* in population *α∈{e, i}*, 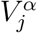 recurrent layer obeys exponential integrate-and-fire (EIF) membrane dynamics^63^:

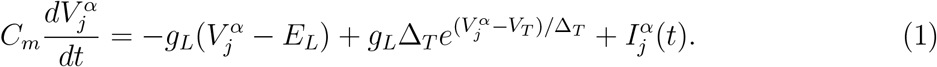

The passive membrane properties are given by the leak conductance *g*_*L*_, the leak reversal potential *E*_*L*_, and the membrane capacitance *C*_*m*_. The second term on the right hand side of Eq.(1) models the excitable membrane nonlinearity that causes an explosive spike onset for 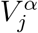above the soft threshold *V*_*T*_ (the sensitivity of spike onset is given by Δ_*T*_). Let time 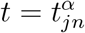 *>*denote the *n*^th^ time that 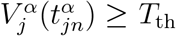 (a threshold crossing), then 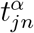 is the *n*^th^ spike time from neuron *j* in population *α*, and we enforced the reset condition 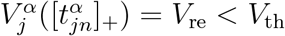.Further, 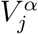 is held constant for a refractory period *τ*_*ref*_ after the reset. The neuron model parameters are set to *τ*_*m*_ = *C*_*m*_*/g*_*L*_ = 15 ms, *E*_*L*_ = *−*60 mV, *V*_*T*_ = *−*50 mV, *V*_th_ = *−*10 mV, Δ_*T*_ = 2 mV, *V*_*re*_ =*−* 65 mV, *τ*_*ref*_ = 1.5 ms. For inhibitory neurons, *τ*_*m*_ = 10 ms, Δ_*T*_ = 0.5 mV, *τ*_*ref*_ = 0.5 ms. Similar model formulations have been used in past studies^11, 27, 64^, and we used the same parameters and constants as in our previous work^27^ unless otherwise specified. There are also model parameters that are not fully explored in previous literature and need to be determined as the goal of the customization. We call them “free parameters”, to be introduced below.

Let the spike train from neuron *k* in population *α ∈ {e, i, F }* be 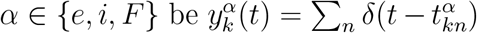, where *d*(*t−s*) is the Dirac delta function centered at time *t* = *s*. The total synaptic current to a neuron *j* in population *α* is:

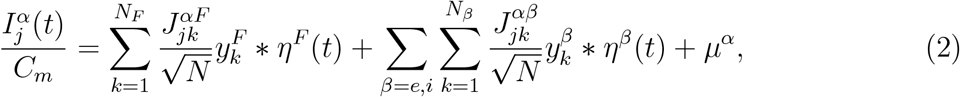

where *N* = *N*_*e*_ +*N*_*i*_ = 3125 and *\** denotes convolution. The synaptic connectivity strength 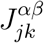 is equal to *J*^*αβ*^ if neuron *k* in population *β* connects to neuron *j* in population *α*, otherwise it is set to zero (*α∈* {*e, i*}and *β∈* {*e, i, F*}). There are then six total synaptic connectivity strengths: *J*^*ei*^, *J*^*ii*^, *J*^*ie*^, *J*^*ee*^, *J*^*eF*^, *J*^*iF*^, which are free parameters. The synaptic kernel from population *β, η*^*β*^(*t*), is given by:

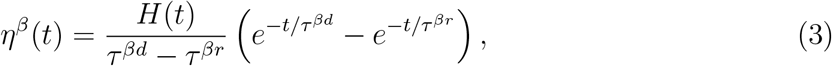

where *H*(*t*) = 0 for *t <* 0 and *H*(*t*) = 1 for *t* ≥ 0. The time constants *τ*^*er*^ = *τ*^*ir*^ = *τ* ^*Fr*^ = 1 ms, *τ* ^*Fd*^ = 5 ms, and *τ*^*ed*^ and *τ*^*id*^ are free parameters.

For the classical balanced network (CBN), the connection probability from a presynaptic neuron in population *β* to a postsynaptic neuron in population *α* is set to a constant 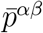. Throughout we used 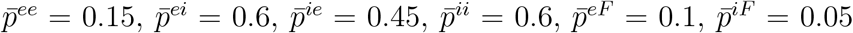, to enable the network to display a similar average number of connections as the model in our previous work^27^.

For the spatial balanced network (SBN), neurons in the two layers are arranged uniformly on a 1 mm square grid. The probability of a connection from presynaptic neuron *k* in population *β* located at position (*x*_*k*_, *y*_*k*_) to postsynaptic neuron *j* in population *α* located at position (*x*_*j*_, *y*_*j*_) is:

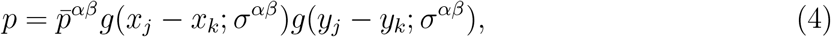

where 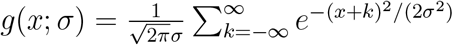 is a wrapped Gaussian distribution. The connection widths (*σ*^*ee*^ = *σ*^*ie*^ = *σ*^*e*^, *σ*^*ei*^ = *σ*^*ii*^ = *σ*^*i*^, *σ*^*eF*^ = *σ*^*iF*^ = *σ*^*F*^) are free parameters for the SBN. Note that mathematically, the SBN is more flexible than the CBN because the latter can be considered a special case of the former: setting the three parameters that control connectivity width to infinity will turn a SBN into a CBN.

The summary of the free parameters common to both the CBN and SBN network models is shown in Table 1. The SBN model has additional free parameters shown in Table 2.

**Table 2:** Free parameters exclusive to the SBN.

| Parameter description | Symbol | Search range |
| --- | --- | --- |
| recurrent inhibitory connection width | $\sigma^i$ | 0 to 0.25 mm |
| recurrent excitatory connection width | $\sigma^e$ | 0 to 0.25 mm |
| feedforward connection width | $\sigma^F$ | 0 to 0.25 mm |

#### Model simulation

To simulate activity from the network, we first instantiated a network model. This involves generating a network connectivity graph based on the connection probabilities, and setting the initial membrane potential of each neuron from a uniform distribution between *−*65 and *−*50 mV, as in our previous work^27^. We will discuss the impact of the randomness induced by the realization of the connectivity graph and initial membrane potentials in the following sections. After model instantiation, the differential equations were solved by the forward Euler method using a time step of 0.05 ms for a duration of the simulation predetermined by the user.

#### Identifying the four activity regimes for the CBN

For Fig. 1 and Fig. 2, we needed to identify CBN parameters that correspond to each of the four activity regimes introduced in Brunel (2000)^34^. Because the network architecture in Brunel (2000) is different from ours (in terms of the number of neurons, choice of integrate-and-fire neuron model, etc.), the parameter values in their paper are not directly applicable to our network. Thus, we randomly sampled 5, 000 parameter sets from the search range of the CBN (Table 1). We then selected parameter sets that produced each of the following four combinations of statistics: low *r*_sc_ and high *ff* (asynchronous irregular), high *r*_sc_ and low *ff* (synchronous regular), high *r*_sc_ and high *ff* (synchronous irregular), and low *r*_sc_ and low *ff* (asynchronous regular). Fig. 1a shows the spike trains of 50 randomly-selected neurons over a period of 200 ms for each activity regime. For the analysis in Fig. 2b, we simulated 140.5 seconds of spiking activity and computed the activity statistics of 50 randomly sampled neurons for each of the four activity regimes (see *Estimating activity statistics)*.

### Activity statistics

Let 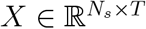 be a matrix of spike counts taken in a fixed time window (defined below) for *N*_*s*_ sampled neurons (either from the neuronal recordings or SNN) and *T* time bins. Based on *X*, we computed the following activity statistics (illustrated in Supplementary Fig. 1):

#### Single-neuron statistics

We considered two commonly used single-neuron statistics: firing rate (*fr*) and Fano factor (*ff*). The *fr* is defined as the mean firing rate across all neurons and trials. Specifically, we average all elements of *X* and divide by the duration of the spike count window.

The *ff* measures the trial-to-trial variability of the activity of each neuron. For each neuron (i.e., row of *X*), we compute its Fano factor as the variance of the *T* values divided by the mean of the *T* values. We then average these Fano factor values across all neurons. For reference, if the spike counts for each neuron were Poisson distributed, then *ff* would equal 1.

#### Pairwise statistic

We considered the pairwise spike count correlation (*r*_sc_), commonly-used to measure how pairs of neurons covary^37^. The *r*_sc_ was computed by first computing the Pearson correlation for each pair of neurons across the *T* trials, then averaging the correlation values across all *N*_*s*_(*N*_*s*_*−*1) pairs of neurons. We applied the Fisher transformation^65^ when comparing *r*_sc_ values in the cost function because it makes the *r*_sc_ values more Gaussian-distributed, as in previous work^66^:

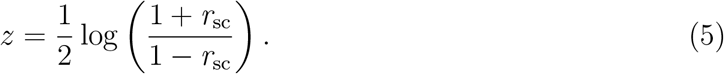

#### Population statistics

We considered three statistics that characterize population-wide covariability: the percent shared variance (%_sh_), the dimensionality of the shared variance (*d*_sh_), and the eigenspectrum of the shared variance (*es*)^31, 32^. These statistics are based on factor analysis (FA), the most basic dimensionality reduction method that partitions variance that is shared among neurons from the variance that is independent to each neuron. Note that principal component analysis (PCA) does not distinguish between these two types of variance.

Using the spike count matrix *X*, we can compute the *N*_*s*_ *× N*_*s*_ covariance matrix *C*. FA performs the decomposition *C≈LL*^*T*^ + Ψ, where 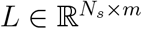 is the loading matrix, 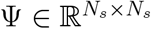 is a diagonal matrix containing the independent variance of each neuron, and *m* is the number of latent dimensions. The matrix *LL*^*T*^ represents the variance shared among neurons (termed the “shared covariance matrix”) and Ψ represents the variance independent to each neuron. The FA parameters *L* and Ψ are estimated from the neuronal activity using the expectation-maximization (EM) algorithm67. The number of latent dimensions *m* is determined by maximizing the 5-fold cross-validated data likelihood.

Based on these FA parameters, we define three population statistics. The %_sh_ quantifies the percentage of each neuron’s variance that is shared with one or more of the other simultaneously-recorded neurons. This value is then averaged across neurons. Specifically, we compute:

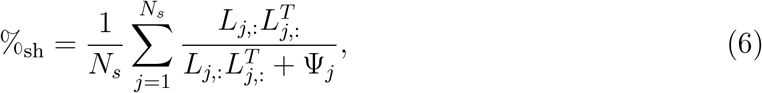

where *L*_*j*,:_ represents the *j*th row of *L*, and Ψ_*j*_ represents the *j*th diagonal element of Ψ. Note that %_sh_ is related, but not equivalent, to *r*_sc_^32^.

The *d*_sh_ measures the complexity of the shared variance among neurons (i.e., the number of population activity patterns needed to describe the shared variance). For example, if all neurons increased and decreased their activity together, *d*_sh_ would equal 1. In principle, we should choose *d*_*sh*_ = *m*. In practice, we first found *m* by maximizing the cross-validated data likelihood, as described above. Then, we chose *d*_sh_ as the number of dimensions needed to explain 95% of the shared variance (based on the eigenspectrum of *LL*^*T*^). This procedure increases the reliability of the estimated dimensionality^31^.

The *es* measures the relative dominance of the dimensions of shared variance. For example, *m* might equal 3, but one dimension might explain far more shared variance than the other two dimensions. Specifically, *es* is defined as the vector of *N*_*s*_ eigenvalues of *LL*^*T*^, where the eigenvalues are ordered from largest to smallest. Only the first *m* eigenvalues are non-zero. We define *es* in this way so that two eigenspectra with different *m* can be directly compared.

#### Estimating activity statistics

To estimate the activity statistics of the SNN with a given parameter set, we first instantiated the model (which includes generating a network connectivity graph and initial membrane potentials, see *Model simulation*). For estimating the activity statistics in Fig. 2, Fig. 4, Fig. 5, and Fig. 6, we repeated this network instantiation procedure 5 times and averaged the estimated statistics over these repetitions to increase estimation reliability (see below). For a given network instantiation, we simulated the network (see *Spiking network models*) to obtain 140.5 seconds of spiking activity. We then removed the first 500 ms of the spike train to ensure the statistics are computed on the spike trains when the network has reached a stable state, similar to Huang et al. (2019)^27^. We also excluded neurons whose firing rate is less than 0.5 spikes per second for stable estimation of variance-based statistics (see *Feasibility constraints*). We then binned the remaining 140 seconds into 700 bins, each of duration 200 ms. We used 140 seconds of activity with 700 bins because empirically such a number of bins is sufficient for a stable estimation of the aforementioned activity statistics while still keeping the simulation time reasonably low (average of 373 seconds, see *Simulation running time*) for our spiking network model with 3, 750 neurons.

For the spike trains corresponding to a given network instantiation, we computed their activity statistics using 50 randomly sampled excitatory neurons in the recurrent layer (similar to Huang et al. (2019)^27^). We sampled 50 neurons to compare model output spike trains directly with that of recorded neuronal population activity, where the number of recorded neurons is typically around 50. We repeatedly sampled 50 neurons without replacement from the network model 10 times. The activity statistics were then computed for each sampled population and averaged across the 10 samplings. This reduces the sampling variance in the estimation of the activity statistics. If there are 5 network instantiations, we further averaged the activity statistics over these instantiations.

For the neuronal recordings, we first excluded neurons with firing rates less than 0.5 spikes per second. We then randomly sampled 50 neurons and 700 trials without replacement for each recording session and condition, where each trial corresponds to a single stimulus presentation (see *Neuronal recordings*). Within each trial, we took spike counts in a 200 ms bin preceding stimulus onset. Hence, the activity statistics for the network model and neuronal recordings are both computed using 50 neurons and 700 trials to ensure consistency for the comparisons.

### Neuronal recordings

Experiments were approved by the Institutional Animal Care and Use Committee of the University of Pittsburgh and were performed in accordance with the United States National Research Council’s Guide for the Care and Use of Laboratory Animals. We reanalyzed data from experiments reported in previous studies^49, 68^. In brief, we trained two rhesus macaque monkeys (monkeys P and W) to perform a spatial attention task. At the beginning of each task trial, the animal first fixated on a central dot for 300–500 ms. Gabor stimuli were presented, one on each side of fixation, for 400 ms. One of the two stimulus locations was block-cued to change its orientation with 90% probability. After the end of the stimulus presentation, a blank inter-stimulus period of 300*−*500 ms followed. The described sequence repeated and on each presentation, there was a fixed probability of one of the Gabor stimuli changing orientation at each presentation (i.e., a flat hazard function). The task of the animal was to detect a change in orientation of one of the two stimuli and make a saccade to the stimulus that changed. Thus, the animal would benefit from maintaining constant attention to the cued location throughout the task trial.

Two 100-electrode Utah arrays (Blackrock Microsystems, one in V4 and one in PFC) were used to record neuronal activity in V4 and PFC simultaneously during the spatial attention task. There were two cue conditions (attention directed to the aggregate V4 receptive field or to the other hemifield) and two stimulus orientations (45° and 135° with the hemifields always containing orthogonal orientations), leading to four unique task conditions with different firing rates and population statistics. For each condition, we included only recording sessions with at least 50 neurons whose firing rate is greater than 0.5 spikes per second each and at least 700 stimulus presentations for accurate estimation of the activity statistics (see *Estimating activity statistics*). This yielded 10 sessions for V4 of monkey W, 20 sessions for PFC of monkey W, 19 sessions for V4 of monkey P, and 19 sessions for PFC of monkey P. More sessions were excluded for V4 than PFC for monkey W because many V4 neurons in monkey W had firing rates less than 0.5 spikes per second. We included both successful and failed trials because we looked at the 200 ms bin preceding stimulus onset which was largely unaffected by the eventual trial result. We customized the network models to each of the four conditions separately (see *Customizing CBN and SBN to neuronal recordings*).

### Cost function

We measured the discrepancy between the SNN-generated activity and neuronal recordings using a cost function. Specifically, our cost function is a weighted linear combination of the normalized distance of each activity statistic from its target value. Let *S* be the set of statistics included in the cost function. For example, *S* = { *fr, ff, r*_sc_, %_sh_, *d*_sh_}, *es* indicates that all six activity statistics are used for customizing the SNN. Let 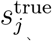 and *s*_*j*_(***θ***) denote the *j*-th activity statistic of the neuronal recordings (i.e., the target value) and that of a network model under parameter set ***θ***, respectively, where *j* ∈ S. The cost function is defined as:

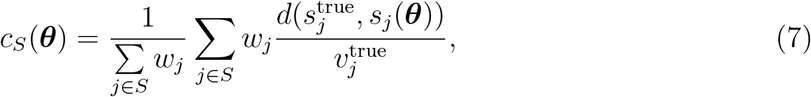

where *d*(·, ·) is a distance function. In this work, 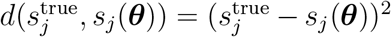. The weight, *w*_*j*_*∈* [0, 1], indicates the relative importance of each statistic and is predefined by the user. If a weight is zero, the corresponding activity statistic is not used during the customization procedure. In this work, we set *w*_*j*_ = 1 for all *j* because we wanted to weigh each statistic equally. The terms 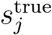 and 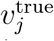 are the mean and variance of the *j*-th activity statistic across simulations or recording sessions (defined below). The variance term serves to down-weight a statistic if its variance is large, indicating the estimation is unreliable. For the eigenspectrum of the shared covariance matrix (*es*), *d*(·,·) is defined as the sum of squared differences of the corresponding elements in the eigenspectra. The variance term for *es* is then computed across sessions or recording sessions using this scalar value.

For the network model (Fig. 4), 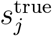 and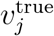 are the mean and variance, respectively, of the corresponding statistic over 5 network instantiations with randomly generated graphs and initial membrane potentials corresponding to the same ground truth parameter set. For neuronal recordings (Fig. 5 and 6), 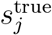 and 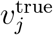 are the mean and variance, respectively, across multiple recording sessions from the same monkey and experimental condition. In the sections below, we will refer to *c*_*S*_(***θ***) as simply *c*(***θ***), where *S* will be clear from the context.

### Optimization algorithm

#### Problem setup

The goal of the optimization algorithm is to find a parameter set ***θ*** *∈* ℝ ^*d*^ in the search region Θ that minimizes *c*(***θ***):

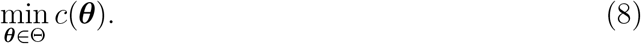

In practice, *c*(***θ***) does not have a closed-form expression in terms of the model parameters and cannot be optimized using gradient-based methods. This is because, for a large-scale spiking network model, *c*(***θ***) depends on several activity statistics, which in turn depend on the computationally demanding numerical simulation of the SNN (see *Simulation running time*). Hence, for a given ***θ***, we cannot compute *c*(***θ***) directly as a function of ***θ***. Instead, we simulate the network to obtain an estimate of *c*(***θ***), denoted *c*ĉ(***θ***). The estimation error is *c*(***θ***) *− ĉ* ĉ(***θ***) and arises from several sources. First, for a given ***θ***, network connectivity graphs are randomly generated. This is because the network connectivity graph is not a parameter of the network model, but is instead drawn from probability distributions specified by the parameters ***θ***. Second, for a given graph, initial membrane potentials of each neuron are drawn randomly to ensure diversity of membrane potentials in the neuronal population, as in our previous work^27^. Third, the network has multiple layers, where the neurons in the first layer (the feedforward layer) emit spikes according to independent Poisson processes. Hence the spike trains from the first layer will differ under the same connectivity graph and initial membrane potentials.

##### Algorithm 1

Random search for SNN customization

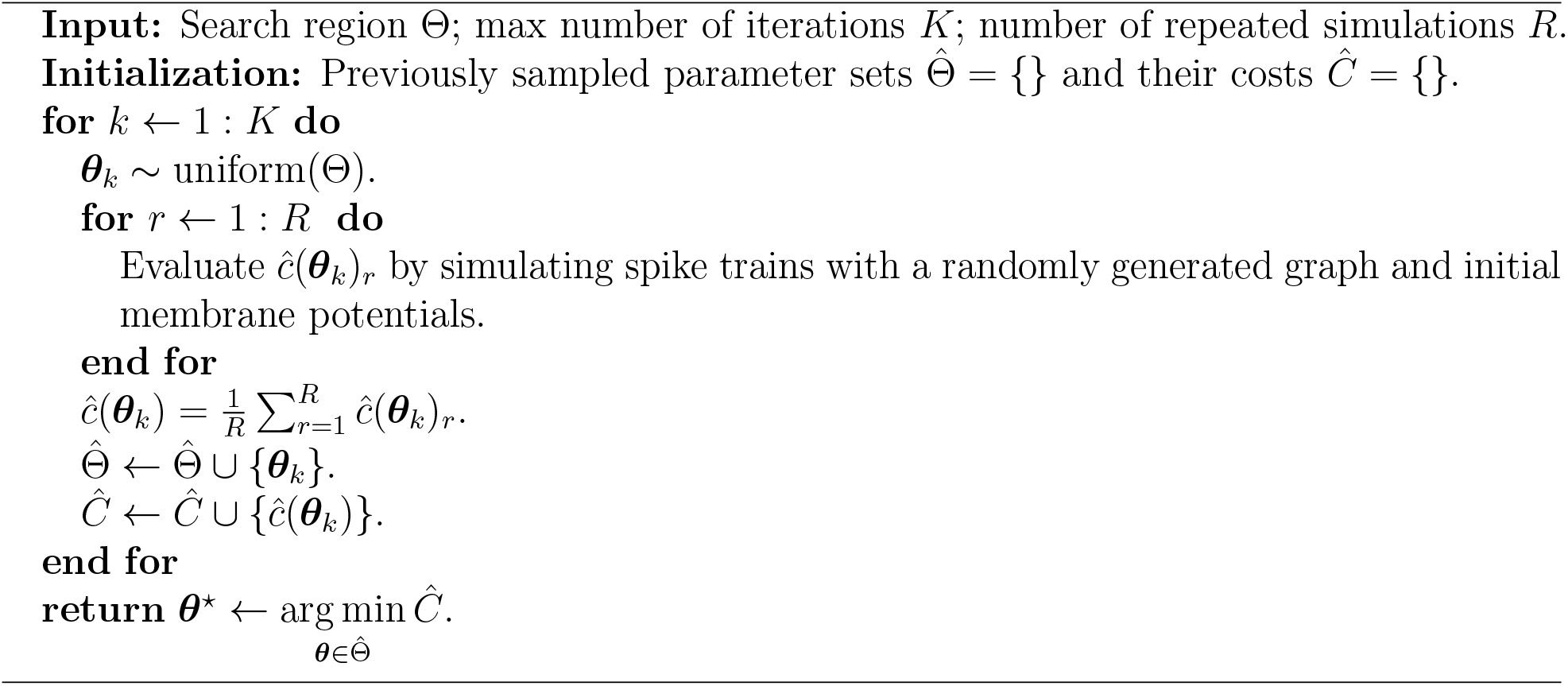

In the following sections, we will introduce two optimization algorithms (Bayesian optimization and random search) to minimize *c*(***θ***), and two innovations (feasibility constraints and intensification) to accelerate optimization. Both innovations can be incorporated into Bayesian optimization or random search. We term Bayesian optimization with both innovations “SNOPS” (Fig. 4, blue), random search with both innovations “accelerated random search” (Fig. 4, red), and random search without innovations “random search” (Fig. 4, green).

#### Random search

An intuitive approach to minimizing *c*(***θ***) without a closed-form expression is random search. Random search is commonly used as a benchmark in optimization and has been shown to have similar performance to more advanced algorithms in many optimization tasks^69^. At each iteration, the algorithm randomly samples a parameter set uniformly from the search region Θ and evaluates its cost. The algorithm terminates after a user-defined number of iterations, *K*, has been reached (Algorithm 1).

To reduce the variance of *ĉ* (***θ***), we repeatedly simulate spike trains with randomly generated graphs and initial membrane potentials using the same parameter set for *R* repetitions (*R* = 5 in this work). For each repetition, we evaluate the cost, then average across the repetitions (Algorithm 1, the inner loop). Note that we will improve this variance-reduction method using one of the innovations (i.e., intensification) to be introduced later.

#### Bayesian optimization

Random search samples parameter sets independently at each iteration and is not guided by the previously-sampled parameter sets. To accelerate the algorithm, we turn to Bayesian optimization (BO). BO utilizes previous evaluations of the cost to guide the parameter search in a way that promotes both exploration and exploitation. BO has been demonstrated to optimize cost functions with fewer iterations than random search in various optimization tasks^44^.

##### Algorithm 2

Bayesian optimization for SNN customization

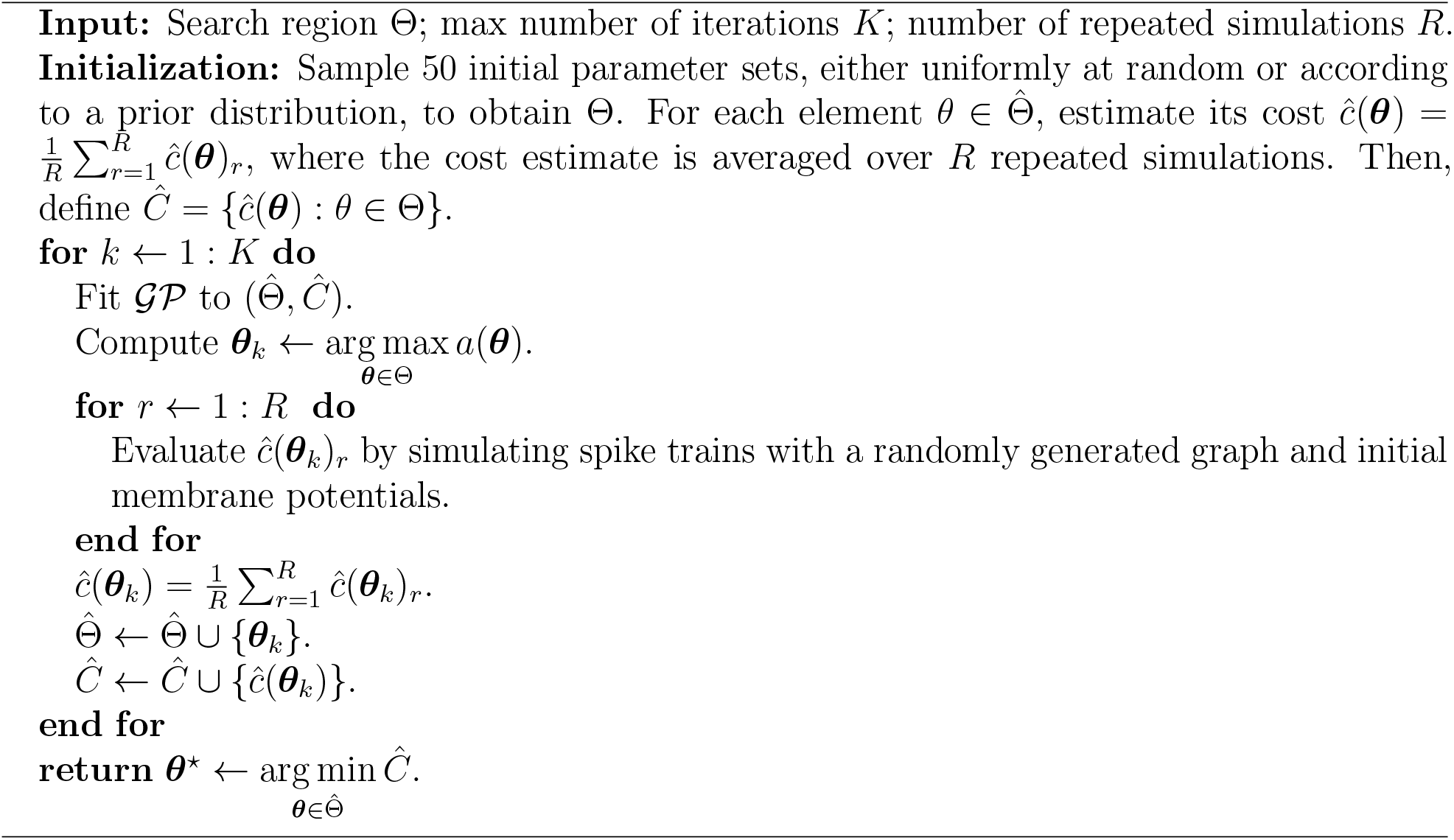

BO involves two major components: (1) a Gaussian process (GP) model^46^ to approximate the cost function and (2) an acquisition function to determine the parameter set to sample at the next iteration. The full algorithm of BO involves iteratively updating the GP model and proposing the next parameter set using the acquisition function. This is outlined in Algorithm 2.

First, BO uses a GP to approximate *c*(***θ***). If two sets of parameters, ***θ***_1_ and ***θ***_2_ are similar, we expect the corresponding costs *c*(***θ***_1_) and *c*(***θ***_2_) to also be similar. To capture this intuition, BO approximates the cost function as a smooth function of the model parameters using a GP. The GP will allow us to predict *c*(***θ***) (posterior mean of the GP) and our uncertainty about the value of *c*(***θ***) (posterior variance of the GP) for a candidate ***θ*** without performing the computationally demanding evaluation of *c*(***θ***) explicitly. Specifically, we write:

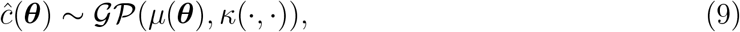

where *µ*(***θ***) : Θ → ℝ is the mean function of the GP and is set as a constant, 0, without loss of generality^46^ . The covariance function, *κ*(·, ·) : Θ *×* Θ → ℝ, is a positive definite kernel function defined on any two points in the search region, Θ. We use the ARD (Automatic Relevance Determination) Matérn 5/2 kernel. The Matérn 5/2 kernel is commonly used in BO because it allows for possible non-smoothness of a cost function. Its ARD variant fits a different length scale for each of the *d* elements of ***θ***, as determined by data. We use the Matlab function fitgpr for fitting a GP model to the sampled parameter sets and their associated costs. In practice, the GP is fitted to the cost estimated by averaging the estimate over *R* = 5 network instantiations of the same parameter set to reduce variance (as in random search). We also log-transformed the cost values when fitting the GP to mitigate the effect of extreme cost values. At the beginning of the optimization, a set of initial parameter sets, Θ, is sampled uniformly to fit the GP since we assume no prior knowledge about the location of the optimal parameter set (Algorithm 2). One may sample Θ according to a prior distribution other than the uniform distribution to guide the initial optimization process if one has such knowledge. We sampled 50 initial parameter sets for Θ, although this number can be varied based on user need. The larger this number, the better the initial GP estimate of *c*(***θ***) will be, but the longer the initialization process will take.

Second, BO uses an acquisition function based on the posterior mean and variance of the GP in Eq.(9) to decide the next parameter set to evaluate. BO selects the parameter set at which the acquisition function value is maximized. This corresponds to a combination of low posterior cost (exploitation, where the cost is predicted to be low) and high posterior variance (exploration, where we have not sampled many parameter sets).

Let 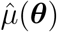denote the posterior mean and 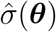denote the posterior standard deviation of the GP at ***θ***. Let *f*_*−*_ be the minimum of *ĉ* (***θ***) over the sampled parameter sets so far. We use the expected improvement^30^ as the acquisition function :

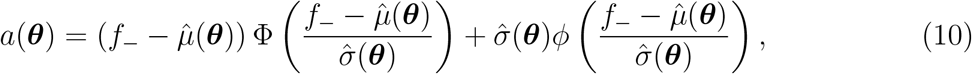

where Φ and *φ* are the normal cdf and pdf, respectively. Eq.(10) is derived based on the goal of preferring ***θ*** whose posterior mean, 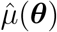, is as small as possible compared to *f*_*−*_ (for the complete derivation, see Brochu et al. (2010)^44^). The first term of the equation represents exploitation: as 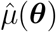 becomes smaller, this term will dominate because 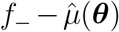 will increase and Φ will approach 1, while *φ* in the second term will approach zero. The second term represents exploration: as 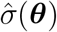 becomes larger, this term will dominate because the first term will approach 0.5 while the second term will increase as *φ* goes towards its peak.

The acquisition function, *a*(***θ***), also does not have an analytical form with respect to ***θ*** because 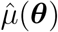 and 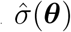 have a non-straightforward dependence on ***θ***. However, 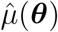 and 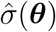 are fast to compute using `fitgpr` in Matlab (typically less than a microsecond for one evaluation). Hence we evaluate *a*(***θ***) on a large number of randomly sampled ***θ*** to quickly maximize *a*(***θ***) (as in the bayesopt function in Matlab). In particular, we first evaluate *a*(***θ***) on 100, 000 randomly sampled parameter sets. We then select 10 parameter sets with the largest *a*(***θ***). We run `fminsearchbnd` to search locally around each of these 10 parameter sets to refine the solution. The final maximizer of *a*(***θ***) is the maximizer from these 10 local searches.

#### Feasibility constraints

Our first innovation seeks to accelerate the optimization process using feasibility constraints. A parameter set, ***θ***, is labeled “infeasible” if, for a particular connectivity graph and initial membrane potentials (a single iteration in the inner loop in Algorithm 1 and 2), it leads to neuronal population activity generated from the network model that falls into either of the following two categories.

First, ***θ*** may lead to extreme firing rates. Low firing rates are undesirable because the resulting spike count matrices are mostly zeros. This can lead to unstable estimates of the variance-based activity statistics (e.g., Fano factor, *r*_sc_, and population statistics). High firing rates are biologically unrealistic (typically *<* 10 spikes/sec for V4 and PFC recordings, see Supplementary Fig. 5). We set the low firing rate threshold as *<* 0.5 spikes/sec and the high firing rate threshold as *>* 60 spikes/sec (mean firing rate across all neurons and time).

Second, ***θ*** may lead to unstable solutions. The network activity may take a period of time to reach a stable state, defined by when the mean firing rate across the neuronal population converges. As noted above, a standard preprocessing step is to remove the first 500 ms of the network-generated spike trains (the period when the network has not yet stabilized)^27^. However, some parameter sets may lead to networks that take more time to stabilize or may never reach the stable state (e.g., switching between multiple stable states). These cases need to be excluded from the customization process because they represent unstable solutions and are not typically used to compare to recorded neuronal activity. Specifically, to determine if a given ***θ*** leads to an unstable solution, we first run change point detection (Matlab `findchangepts` function)^70^ on the time course of the population-averaged mean firing rate after removing the first 500 ms. We then deem ***θ*** to be infeasible if the mean firing rate (across neurons and time) before the change point and that after the change point exceeds a threshold. The threshold is computed as three standard deviations of the mean firing rate (across neurons and time) after the change point.

To speed up the customization process, we wish to rule out infeasible parameter sets with minimal computation. We propose to use a “freeze-thaw” method^71^ by first running a short simulation to generate 10 seconds of spike trains to estimate the feasibility of a parameter set and only proceed to the full simulation (140 seconds, see *Estimating activity statistics*) if the parameter set is feasible. We use 10 seconds for the short simulation because the two constraints only depend on the firing rate, and empirically the estimation of the firing rate tends to stabilize within 10 seconds. However, estimating population statistics, for example, %_sh_, requires substantially more simulation time hence a full simulation is still needed to compute all activity statistics.

For random search, feasibility constraints can be incorporated in a straightforward manner: for each sampled parameter set, if feasible, the algorithm will run the full simulation of 140 seconds to compute the cost. If the sampled parameter set is infeasible, it will simply proceed to the next sampled parameter set. Note that feasibility is evaluated at each of the *R* repetitions in the inner loop (Algorithm 1). The inner loop aborts as soon as the parameter set is deemed infeasible.

For BO, we incorporated the feasibility constraint into the optimization problem^72^:

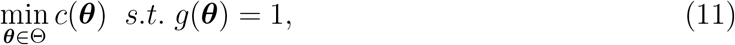

where *g*(***θ***) = 1 if ***θ*** is feasible and *g*(***θ***) = 0 otherwise.

To incorporate this constraint, two parts of BO will change. First, in addition to the GP that approximates the cost function (the GP in Eq.(9)), there is a separate GP that represents the feasibility function, *g*(***θ***), which is fitted to the sampled parameter sets and their feasibility values (which are binary). Second, the acquisition function will now incorporate the GP for the feasibility function. Let 𝒢 𝒫_*c*_ represent the GP on *c*(***θ***) as in Eq.(9), and 𝒢 𝒫 _*g*_ represent the GP on *g*(***θ***). Let 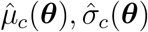 denote the posterior mean and standard deviation, respectively, of 𝒢 𝒫 _*c*_. Similar to Eq.(10), *f*_*−*_ is the minimum of 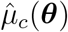 over the parameter sets evaluated so far. The expected improvement (i.e., acquisition) function for the constrained Bayesian optimization becomes^72^:

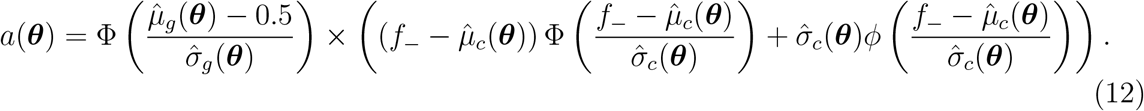

This differs from Eq.(10) in the term Φ 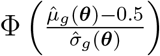 which yields a larger acquisition value if the feasibility posterior mean is high. Note that even though 𝒢 𝒫_*g*_ is fitted to binary values (*g*(***θ***) is either 0 or 1), its posterior mean, 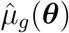, is continuous-valued. Furthermore, in this term, 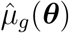 is referenced to 0.5 to ensure symmetry.

#### Intensification

The second innovation seeks to improve the accuracy of estimating the cost with less time. The estimation error arises from several sources, described in *Problem setup*. A high estimation error may result in a final parameter set returned by the algorithm with a low cost in a single evaluation, but with a higher cost if evaluated and averaged over multiple repetitions^73^.

One possible solution is to repeatedly run simulations under the same parameter set for *R* repetitions, as in Algorithm 1 and 2. However, this can be computationally demanding because each repetition corresponds to a lengthy simulation. To avoid performing *R* repetitions for every ***θ*** sampled, we propose to use an intensification algorithm^73^. The main idea is to only perform *R* repetitions of the simulation if we encounter a potentially optimal parameter set. We first define the incumbent parameter set as the parameter set whose cost, calculated by averaging over *R* repetitions, is the smallest over the list of sampled parameters. Similarly, the incumbent cost and standard deviation are the associated mean and standard deviation of the cost of the incumbent parameter set over the *R* repetitions. A sampled ***θ*** is considered potentially optimal if its cost evaluated in the first repetition is within one standard deviation of the incumbent cost. Otherwise, we include it in the list of sampled parameters and proceed to sample the parameter set for the next iteration. The algorithm will perform evaluations for *R* repetitions only for the potential optimal parameter sets. If its average cost over the *R* repetitions is smaller than the incumbent cost, ***θ*** becomes the incumbent parameter set. Otherwise, we still include it in the list of sampled parameters and its associated cost value is the average over the *R* repetitions.

We adopt this method and introduce an additional stopping criterion: the algorithm will stop performing repetitions if the standard deviation of the cost across the performed repetitions is below a specified threshold. A small standard deviation indicates the estimate of the cost of this parameter set is consistent across repetitions and needs no variance reduction. Note that at least two repetitions need to be performed to compute the standard deviation, and if this stopping criterion is met, we follow the same procedure above in determining if ***θ*** becomes the incumbent parameter set. The additional stopping criterion further reduces the total number of simulations throughout the optimization procedure and provides additional acceleration. We set the predefined number of repetitions to *R* = 5 and the standard deviation threshold to 0.15. A larger predefined number of repetitions and a smaller standard deviation threshold will yield a smaller estimation error, at the expense of greater simulation time.

#### Local optima

If SNOPS is run for infinite time, it is guaranteed to return the global optimal parameter set^44, 74^. In practice, finite running time may result in the algorithm returning a local optimum. Empirically, we verified that SNOPS reliably returned a set of activity statistics that matched the recorded neuronal activity when the optimization algorithm was initialized with different initial parameter values (Supplementary Fig. 5). This indicates that, even if the algorithm returns a local optimum, this optimum corresponds to activity statistics that match those of the recorded activity.

### Trade-off cost

We define a trade-off cost to measure whether more accurately reproducing one activity statistic leads to less accurately reproducing another activity statistic (Fig. 6). Intuitively, customizing two statistics *s*_*a*_ and *s*_*b*_ simultaneously might incur a larger cost (of *s*_*a*_ and *s*_*b*_) than customizing each of them individually. The gap between the cost of customizing *s*_*a*_ and *s*_*b*_ together versus individually represents how much the two statistics trade-off with each other. Note that a similar idea has also been explored to show the trade-off in the loss function of a model trained to perform two tasks simultaneously^75^.

For two statistics *s*_*a*_ and *s*_*b*_, let 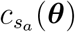 and 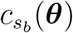 represent the cost values of the optimal parameter sets when individually customizing *s*_*a*_ and *s*_*b*_, respectively. Let 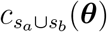 represent the cost resulting from customizing the two statistics together. The trade-off cost between *s*_*a*_ and *s*_*b*_ is defined as:

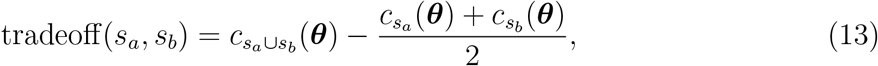

where the second term represents the average of 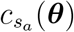 and 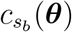. This average makes the second term comparable to the first term.

The trade-off cost is guaranteed to be non-negative because customizing both statistics simultaneously is as challenging or more challenging than customizing each of them separately. This leads to a higher cost for each statistic when the model is customized to both statistics together (first term) as compared to when the model is customized to each of them separately (second term). A trade-off cost of zero indicates that the ability of the model to reproduce one statistic is unaffected by the incorporation of another statistic into the cost function.

### Verifying SNOPS in simulation

To validate the performance of SNOPS in simulation (Fig. 4, Supplementary Figs. 3 and 4), we customized network models to the activity generated from the same type of model. We first randomly sampled 100 parameter sets from the search region (see *Spiking network models*) for the CBN and SBN. We estimated the activity statistics for each sampled parameter set over 5 network instantiations (see *Cost function*). We excluded parameter sets resulting in *fr* smaller than 1, *d*_sh_ smaller than 1, and *ff* larger than 5 because they do not fall within the range of the V4 and PFC activity statistics (Supplementary Fig. 8). We then randomly sampled 40 of the remaining parameter sets for both the CBN and SBN. We chose to use 40 parameter sets because they represent a diverse combination of activity statistics while having reasonable running time. We applied SNOPS, random search, and accelerated random search to customize the CBN and SBN separately. We ran each optimization-based method (SNOPS, random search, and its accelerated variant) for 168 hours (7 days). We found the cost usually plateaus within 168 hours, indicating that the running time is sufficient for SNOPS to converge (Supplementary Fig. 3).

### Customizing CBN and SBN to neuronal recordings

To compare the CBN and SBN as well as to validate the performance of SNOPS on neuronal recordings (Fig. 5, Supplementary Figs. 3 and 4), we ran SNOPS, accelerated random search, random search, and SNPE on the 16 datasets comprising two monkeys (monkeys P and W), four conditions (two cues by two saccade locations), and two brain areas (V4 and PFC). As in the previous section, we set the stopping criterion for the optimization-based methods to 168 hours (7 days) because we found the cost usually plateaus within 168 hours (Supplementary Fig. 3).

To compare the trade-offs of different subsets of statistics between the CBN and SBN (Fig. 6), we customized each network model to the example macaque V4 dataset in Fig. 2 with different subsets of activity statistics included in the cost function. For each customization run, we set the weight of each statistic (*w*_*j*_ in Eq.(7)) to be either 0 or 1, leading to a total of 2^6^*−*1 = 63 customization runs for each network, which accounts for all possible subsets of activity statistics. For each customization run, we ran SNOPS for 168 hours.

### Simulation running time

The running times indicated below were obtained using cluster machines with 40 Intel Xeon Gold 6230 2.10 GHz CPU cores and 250 GB of RAM. For clarity, here we refer to one *customization run* as the process of customizing a SNN to one dataset using SNOPS (Algorithm 2). We used one CPU core for each customization run. The overall running time for one customization run is 168 hours, corresponding to 1, 200 optimization iterations on average. Each optimization iteration involves the following components. First, for a selected parameter set that maximizes the acquisition function, we randomly instantiated a network connectivity graph and initial membrane potentials and generated spike trains from the spiking network with 3, 750 neurons. This is the most time-consuming part of each iteration. It takes 23 seconds to generate 10 seconds of spike trains to determine feasibility (see *Feasibility constraints*) and 373 seconds to generate 140 seconds of spike trains. The values are the same for the CBN and SBN since they have the same number of neurons (3, 750). Second, for the generated spike trains of 140 seconds, it takes 69 seconds to compute its activity statistics. Finally, it takes 32 seconds to select the parameter set for the next iteration, including fitting the GP for both the feasibility constraints and cost function, as well as maximizing the acquisition function. Note that, for some iterations, the spike train generation and activity statistics computation may be repeated up to 5 times due to the intensification procedure (see *Intensification*).

## Data Availability

The V4 and PFC recordings we analyzed for SNN customization are available at the following link: https://kilthub.cmu.edu/articles/dataset/Utah_array_recordings_from_visual_cortical_area_V4_and_prefrontal_cortex_with_simultaneous_EEG/19248827.

## Code Availability

Matlab code for the SNOPS algorithm will be made publicly available upon publication.

## Acknowledgements

This work was supported by NIH R01 NS121913 (C.H.), NIH K99 EY025768 (A.C.S.), NIH R01 EB026953 (B.D., M.A.S. and B.M.Y.), NIH R01 MH118929 (B.M.Y. and M.A.S.), NSF NCS BCS 1734916/1954107 (B.M.Y. and M.A.S.), Simons Foundation NC-GB-CULM-00002794-06 (C.H.), 542967 (B.D.), and 543065 (B.M.Y.), NSF NCS DRL 2124066 (B.M.Y. and M.A.S.), NIH R01 EY029250 (M.A.S.), NIH U19 NS107613 (B.D.), Vannevar Bush faculty fellowship N00014-18-1-2002 (B.D.), NSF NCS BCS 1533672 (B.M.Y.), NIH R01 HD071686 (B.M.Y.), NIH R01 NS105318 (B.M.Y.), NIH RF1 NS127107 (B.M.Y.), and NIH R01 NS129584 (B.M.Y.). This work used the Extreme Science and Engineering Discovery Environment (XSEDE), which is supported by National Science Foundation grant ACI-1548562.

---

**Supplementary Figure 1:**
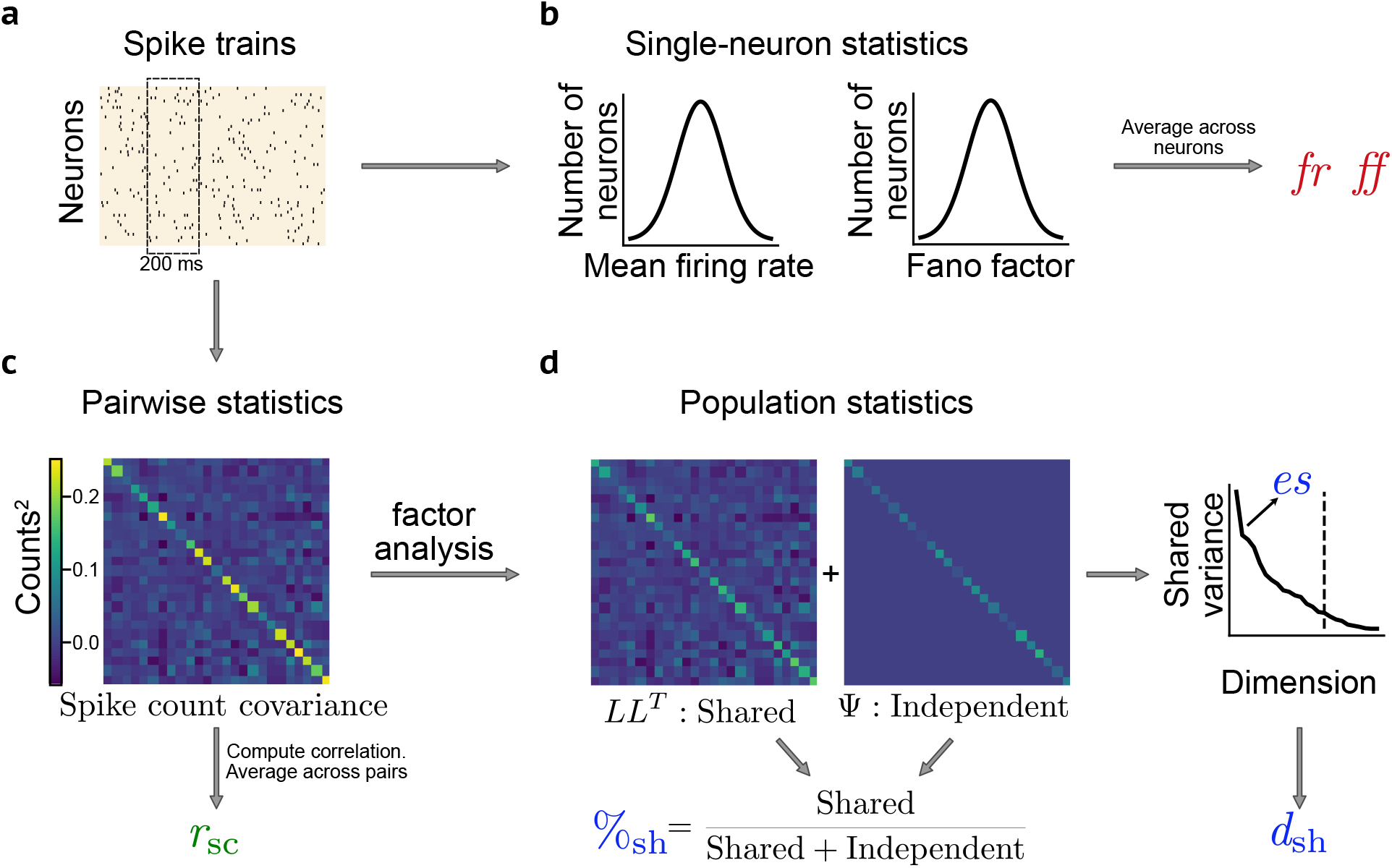
Computing single-neuron, pairwise, and population activity statistics. We introduced three types of activity statistics in Fig. 2. Here we illustrate how to compute these statistics from neuronal activity. **a**, The spike count of each neuron is taken within a specified time window, yielding a *N*_*s*_*×*1 spike count vector for *N*_*s*_ neurons. We can obtain many such spike count vectors taken in non-overlapping time windows, resulting in a *N*_*s*_*×T* spike count matrix, with *T* being the number of non-overlapping time windows. **b**, For the single-neuron statistics, the mean firing rate (*fr*) is computed by averaging the spike counts across time and neurons and dividing them by the duration of the spike count window. The Fano factor (*ff*) is computed as the ratio of the variance of the counts divided by the mean of the counts for each neuron, then averaged across neurons. **c**, The spike count correlation (*r*_sc_) between pairs of neurons is obtained by first computing the *N*_*s*_*×N*_*s*_ covariance matrix of spike counts. The Pearson correlation is computed for each pair of neurons, then averaged across pairs. **d**, For the population statistics, the spike count covariance matrix is first decomposed into a shared covariance matrix (*LL*^T^) and an independent variance matrix (Ψ) using factor analysis^31^. The percent shared variance (%_sh_) is the percent of each neuron’s spike count variance that is shared with other neurons in the recorded population. This value is then averaged across neurons. The dimensionality (*d*_sh_) is the number of latent dimensions needed to explain 95% of the shared variance. The eigenspectrum (*es*) comprises the amount of shared variance represented by each latent dimension. Mathematically, it is the eigenspectrum of *LL*^T^.

**Supplementary Figure 2:**
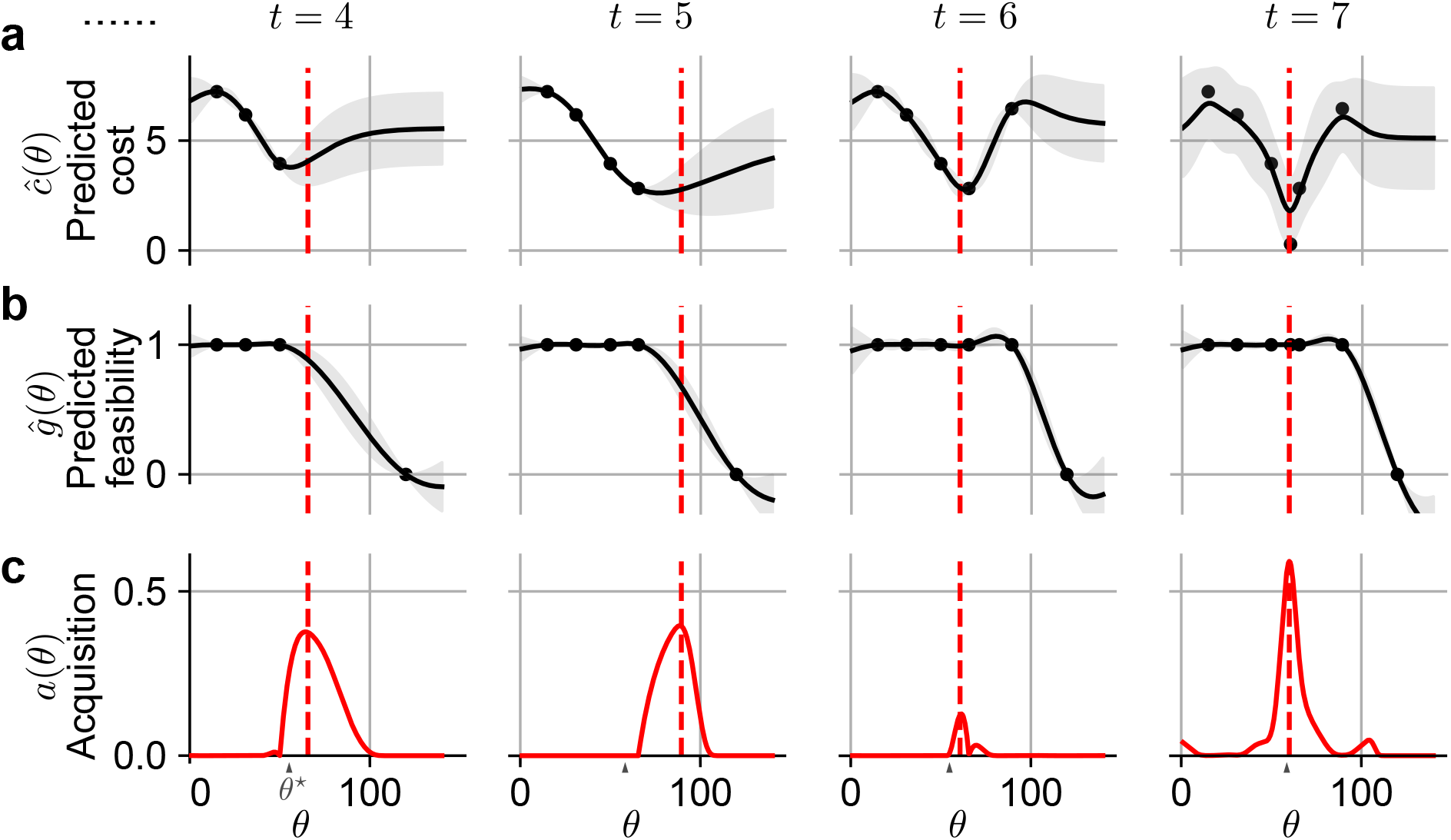
Using SNOPS to customize a SNN with one parameter. We provided an overview of Bayesian optimization in SNOPS in Fig. 3. Here we demonstrate the steps in more detail, in particular those shown in Figs. 3d and 3e. For illustrative purposes, we consider the optimization of a single model parameter *θ*. Details: We first generated spike trains from an SBN whose connection strength, *J*^*eF*^ (see Methods), was set to 60 mV. The values of the rest of the model parameters are set as: *τ*^*id*^ = 8 ms, *τ*^*ed*^ = 5 ms, *J*^*ei*^ =*−*60 mV, *J*^*ie*^ = 10 mV, *J*^*ii*^ =*−*75 mV, *J*^*ee*^ = 20 mV, *J*^*iF*^ = 25 mV, *σ*^*i*^ = 0.1 mm, *σ*^*e*^ = 0.1 mm. The goal of SNOPS here is to customize a separate SBN to these generated spike trains, where *J*^*eF*^ is the only model parameter that is unknown. Because this illustration applies to any model parameter, we denote *J*^*eF*^ as *θ* and the ground truth value of *J*^*eF*^ as *θ*^*\**^. **a**, A Gaussian process (GP) is used to approximate the cost function, (*θ*). Same conventions as in Fig. 3d. With each iteration of SNOPS (left to right), we evaluate the cost at one additional value of *θ* (dashed red line). This leads to an update of both the mean (black line) and uncertainty (gray shading) of the GP, which represents an interpolation of the evaluated costs. Note that the black line might not necessarily pass through all the dots due to the smoothness assumption of the GP, where the smoothing is determined by the Matérn 5/2 kernel (see Methods). **b**, A separate GP represents the feasibility function, ĝ(*θ*). For example, a value of *θ* would be infeasible if it leads to an unrealistically high firing rate (see *Feasibility constraints*). The rationale for using a GP is that, if a value of *θ* is feasible (or infeasible), similar values of *θ* are also likely to be feasible (or infeasible). A parameter *θ* is feasible if ĝ (*θ*) = 1 and infeasible if ĝ(*θ*) = 0. As in **a**, the GP is updated with each iteration of SNOPS (left to right) based on whether the evaluated *θ* is feasible or infeasible (black dots). Note that, for *t* = 4, there are three dots in **a**, but four dots here in **b**. The reason is that the rightmost dot (representing the iteration before *t* = 4) corresponds to an infeasible value of *θ* (i.e., *ĝ* (*θ*) = 0), and hence the cost at that *θ* was not evaluated. **c**, Based on the predicted cost function (from **a**) and feasibility function (from **b**), SNOPS computes an acquisition function, *a*(*θ*). The maximum of the acquisition function (dashed red line) determines the *θ* to evaluate during the next iteration of SNOPS. Intuitively, the acquisition function identifies values of *θ* that are feasible and for which the predicted cost is low. In addition, the acquisition function implements an exploration-exploitation trade-off, whereby values of *θ* for which the cost is uncertain are favored. As SNOPS iterates (left to right), the maximum of *a*(*θ*) approaches the ground truth value *θ*^*\**^.

**Supplementary Figure 3:**
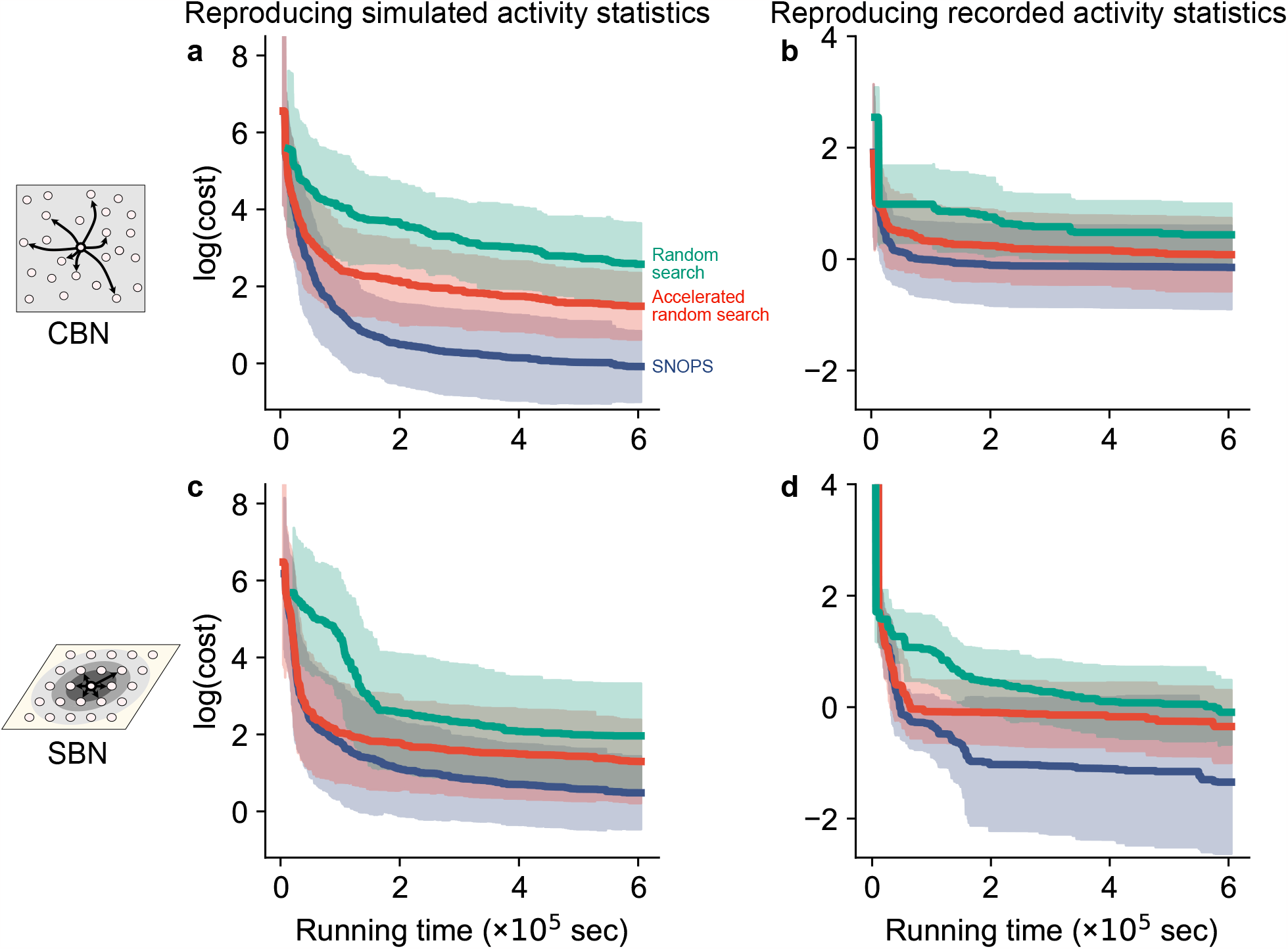
SNOPS outperforms random search methods on simulated activity and neuronal recordings. In Fig. 4, we showed that SNOPS outperforms random search methods when customizing a CBN to simulated activity. Here we compare SNOPS, random search, and accelerated random search in a wider range of settings, involving also SBN and neuronal recordings. Same conventions as Fig. 4b. **a**, Customizing a CBN to simulated activity. These curves are identical to those shown in Fig. 4b. **b**, Customizing a CBN to neuronal recordings in macaque V4 and PFC. These curves correspond to the results for the CBN in Fig. 5c. **c**, Customizing a SBN to simulated activity. **d**, Customizing a SBN to neuronal recordings in macaque V4 and PFC. These curves correspond to the results for SBN in Fig. 5c. Each curve indicates how the cost (which we seek to minimize, plotted in log scale) changes during the optimization procedure, as a function of the computer running time. The shading represents mean *±*1 SD across the customization runs. Panels **a** and **c** include 40 customization runs, as in Fig. 4. Panels **b** and **d** include 16 customization runs corresponding to the neuronal recordings (2 monkeys *×* 2 brain areas *×* 4 experimental conditions) in Fig. 5c. The cost decreases with running time for all three algorithms on both simulated activity and neuronal recordings, as expected. The cost plateaus for all algorithms, suggesting that the stopping criterion we set (see Methods) is sufficient for the algorithms to converge. We found that SNOPS outperforms the random search methods for both simulated data and recorded activity for both CBN and SBN (the blue curves are below the red and green curves). This result demonstrates that Bayesian optimization enables SNOPS to find a better solution than random search methods for a given amount of computer running time. Accelerated random search also outperforms random search across datasets and models (the red curves are below the green curves), suggesting that the two innovations (feasibility constraint and intensification, see Methods) further accelerate the customization process. The amount by which Bayesian optimization outperforms the random search methods depends on the specific network model and the activity to which the model is being customized. For example, SNOPS performed similarly to accelerated random search in **b**. This likely occurs because the CBN is unable to generate a wide range of population statistics, and so there are many parameter sets that correspond to a similar minimal cost. As a result, the random search methods can readily find one of these parameter sets. Even so, SNOPS performed as well or better than the other two methods.

**Supplementary Figure 4:**
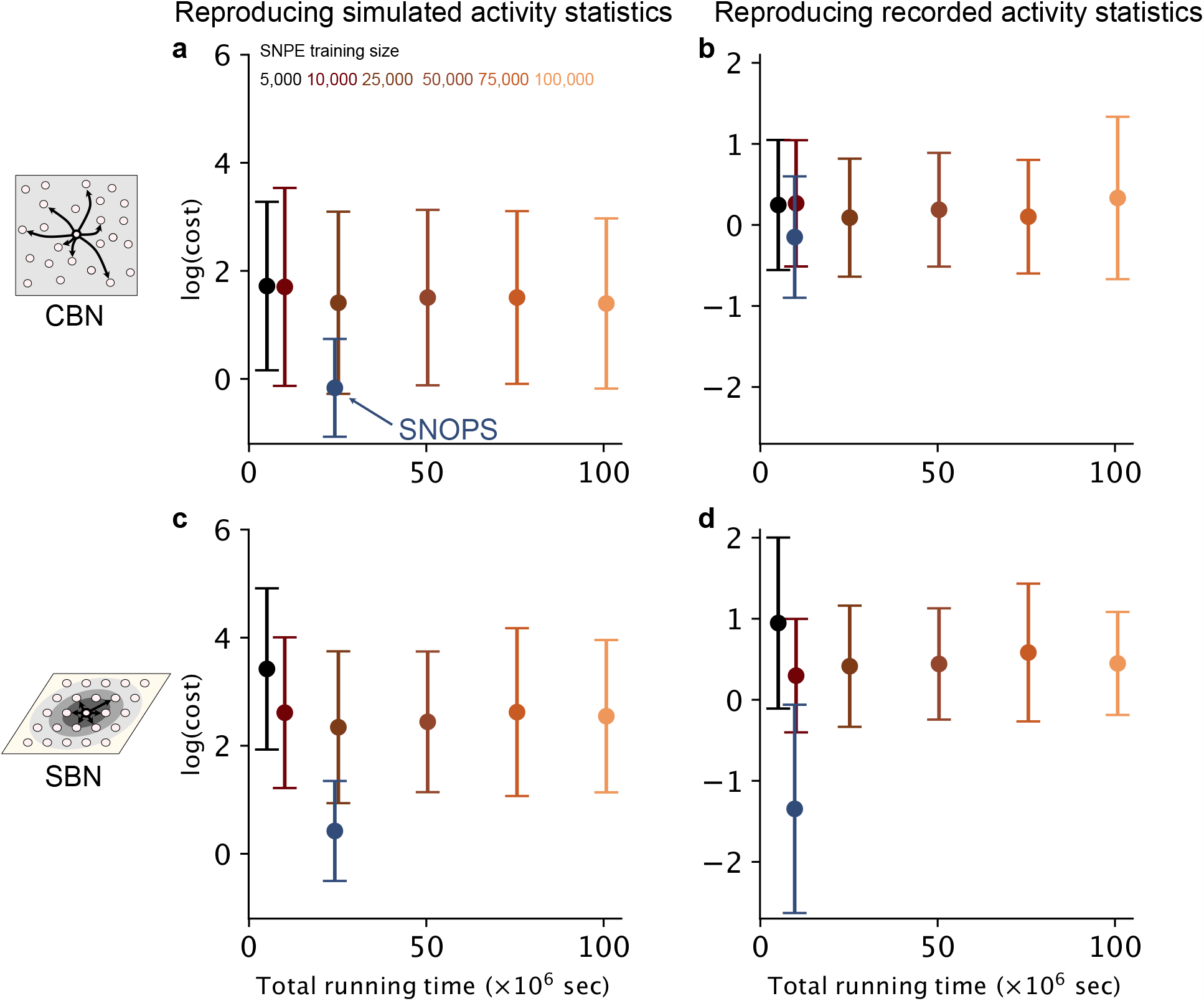
SNOPS outperforms SNPE on simulated activity and neuronal recordings. In Fig. 4 and Supplementary Fig. 3, we showed that SNOPS outperformed random search methods when customizing both CBN and SBN to simulated and recorded neuronal activity. Here we compare SNOPS to a method that uses deep-learning-based inference, Sequential Neural Posterior Estimation (SNPE)^29^, using the same models and neuronal recordings as in Supplementary Fig. 3. One might also consider using Emergent Property Inference (EPI)^52^. However, EPI is not applicable to large-scale SNNs because it requires the cost function to be differentiable with respect to the model parameters. SNPE uses deep neural networks to approximate the distribution of the model parameters given the recorded activity statistics. Applying SNPE involves a training stage and an inference stage. During the training stage, a deep generative model is trained on a large number of simulations comprising the ground truth parameter set of each simulation and its corresponding activity statistics. During the inference stage, given the recorded activity statistics, SNPE returns samples of parameter sets from a posterior distribution representing the likelihood of the parameters consistent with the recorded activity. We wish to compare the two methods in terms of their optimal cost and total running time. There are two key differences between SNOPS and SNPE: (1) SNOPS returns a single parameter set whereas SNPE returns a distribution of parameter sets. The choice between these two outputs depends on the number of customized models desired for the scientific goal (see Discussion). (2) SNPE requires a large number of computationally demanding simulations for its training stage. It can then perform fast parameter inference on any new dataset without the need to run additional simulations. In contrast, SNOPS does not need simulations upfront. However, for each new dataset, SNOPS requires a separate customization run consisting of computationally demanding simulations in the iterative procedure. Comparing the two methods fairly is challenging because they are tailored for different use cases (see Discussion). We did the following to ensure fairness to the best of our abilities (details included below): (1) To compare the parameter distribution inferred by SNPE to a single parameter set returned by SNOPS, we used the mode of the SNPE distribution as the optimal parameter set for SNPE. (2) For the total running time of SNOPS, we considered the time required for a single customization run, which is equivalent to one dataset, and multiplied it by the number of datasets. Each customization run takes 168 hours to converge, as illustrated in Figure 3. Therefore, the total running time for SNOPS would be 40*×*168 hours for the simulated datasets and 16*×*168 hours for the recorded datasets. For SNPE, the total running time includes two parts. The first part consists of generating simulation samples of a given number (training size) and training deep networks, which is the same for both simulated and recorded activity. We systematically varied the training size from 5, 000 to 100, 000 simulations to examine its impact on the fitting quality of SNPE. The second part is the inference time for the 40 simulated datasets (panels **a** and **c**) or the 16 recorded datasets (panels **b** and **d**). For both methods, all other settings, including the size of the network model and the datasets to fit, were identical. The black-to-orange dots represent the cost of SNPE (in log scale) under different training sizes. The blue dots represent the cost and total running time of SNOPS. The error bars represent mean *±*1 SD across the datasets. Panels **a** and **c** include 40 simulated datasets generated by CBNs or SBNs, as in Supplementary Fig. 3a and c. Panels **b** and **d** include 16 recorded datasets (2 monkeys *×*2 brain areas *×*4 experimental conditions) as in Supplementary Fig. 3b and d. We found that SNOPS outperforms SNPE: SNOPS (dark blue dot) has a lower cost than SNPE (black-to-orange dots) for different SNPE training sizes. The main reason is that SNPE requires a large number of computationally demanding simulations to ensure that the deep neural networks can accurately characterize the relationship between model parameters and activity statistics. SNPE may need more than 100, 000 training samples to achieve comparable results to SNOPS. However, the high computational cost for SNPE can be justified by the benefits of a richer characterization of the parameter space since it returns a distribution of parameter sets and provides fast inference for new datasets. Therefore, the choice between SNOPS and SNPE depends on the network simulation time and the goal of model customization. When the simulation time is low, SNPE can be used to obtain a parameter distribution and make fast inferences on a large number of new datasets. In this case, SNOPS may incur a greater running time because SNOPS starts from scratch for each new dataset. When the simulation time of the model is substantial (as for the CBN and SBN), generating a sufficient amount of training samples for SNPE can be computationally prohibitive. In this case, one can instead use SNOPS to obtain a single optimal parameter set for a given dataset. Details: For SNOPS, the optimal cost for each dataset (simulated or recorded) is the result of one customization run (Supplementary Fig. 3, cost values of the right end of the blue curves). For SNPE, we trained the deep generative model using different numbers of network simulations (training sizes): 5, 000, 10, 000, 25, 000, 50, 000, 75, 000, and 100, 000. For each training size, we first sampled the corresponding number of parameter sets according to a uniform distribution, which is the default setting in SNPE. For each sampled parameter set, we generated spike trains under one connectivity graph and computed its activity statistics. We then fed the sampled parameter sets, as well as their activity statistics, to the SNPE training algorithm and ran it until convergence. We did not include training sizes larger than 100, 000 because it would require over 25, 000 CPU hours, which was computationally prohibitive. At the inference stage, for a set of activity statistics corresponding to a simulated or recorded dataset, we drew 10, 000 parameter sets from the SNPE distribution. We selected the single optimal parameter set with the largest log-likelihood as the optimal parameter set returned by SNPE, representing the mode of the posterior distribution. We computed the optimal cost for SNPE using five network instantiations of connectivity graphs and initial membrane potentials corresponding to the same identified parameter set in the same way as in SNOPS.

**Supplementary Figure 5:**
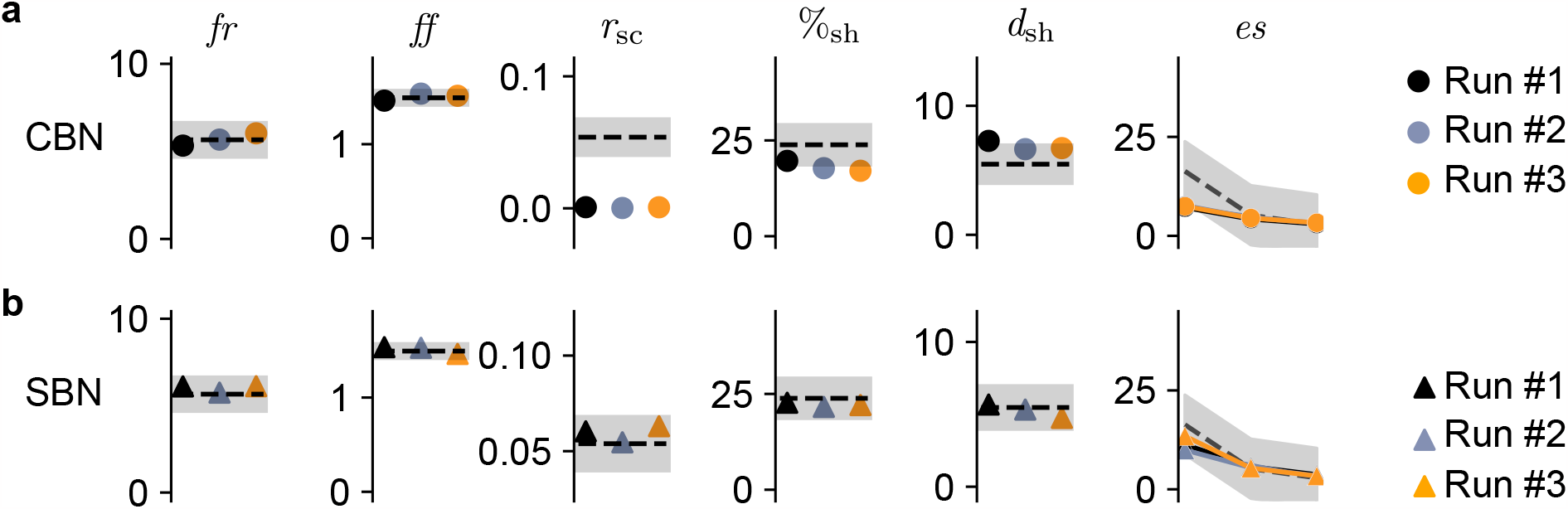
SNOPS returns consistent results across multiple runs. In Fig. 5, we compared the performance of the CBN and SBN in reproducing the activity statistics of recordings in area V4 and PFC. Here we test the reliability of the results returned by SNOPS. Ideally, we would assess this reliability by ensuring that it returns the “ground truth” parameters for each recording. However, because the real neuronal recordings were not generated by a model, there are no “ground truth” parameters. Instead, we assessed whether varying the initialization of SNOPS would affect results. There were several initialization settings that could affect the outcome of the customization process: the initially sampled seed parameter sets for SNOPS (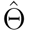in Algorithm 2), the randomly generated connectivity graph, the randomly configured initial membrane potentials for each neuron, and the stochasticity of the activity in the feedforward layer (see Methods). To investigate if SNOPS returned consistent results, we applied SNOPS to customize CBNs (circles) and SBNs (triangles) to the same V4 dataset as in Fig. 5a and 5b for three customization runs (three colors). For each customization run, the settings of SNOPS (initially sampled parameters, connectivity graph, initial membrane potentials, activity in the feedforward layer) were randomized (see Methods for the ranges of the values). We found that, regardless of the initialization, SNOPS returned highly reliable results across the three runs (three dots have similar values for all panels). The costs of the customized parameters were also similar across the three runs: 2.72, 2.72, 2.74 for the CBN; 0.24, 0.26, 0.24 for the SBN. Hence, SNOPS is robust to different initializations of the optimization procedure.

**Supplementary Figure 6:**
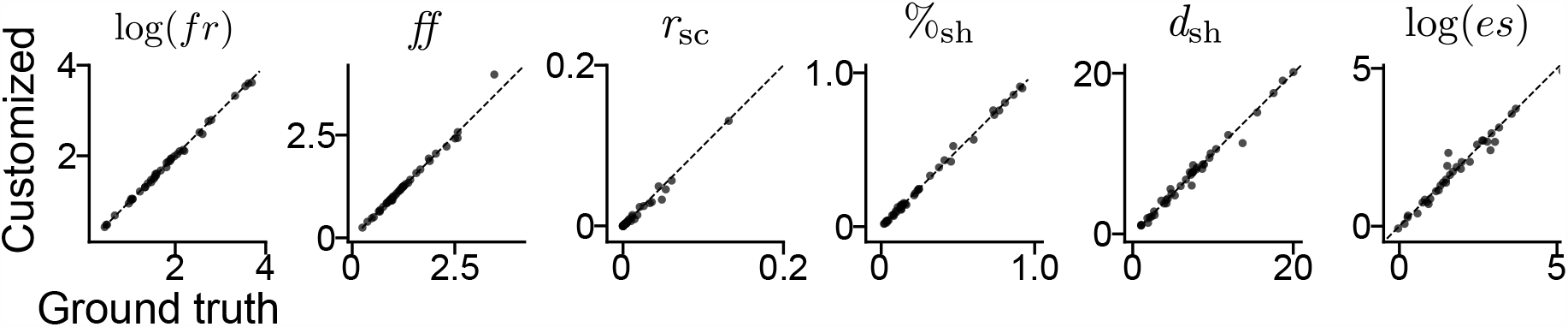
SBNs accurately recover ground truth activity statistics. In Fig. 4, we validated the performance of SNOPS in customizing the CBN to simulated activity. Here we perform the same analysis with the SBN. As we did with the CBN, we randomly drew 40 parameter sets for the SBN and used them to simulate spiking activity. We then used SNOPS to customize a SBN to the simulated activity (see *Verifying SNOPS in simulation*). Same conventions as in Fig. 4d. We found that SNOPS was able to find model parameters for the SBN that accurately reproduced the activity statistics (all dots lie near the diagonal line). For *r*_sc_, one data point (0.65, 0.68) fell outside of the plotting range and is not shown.

**Supplementary Figure 7:**
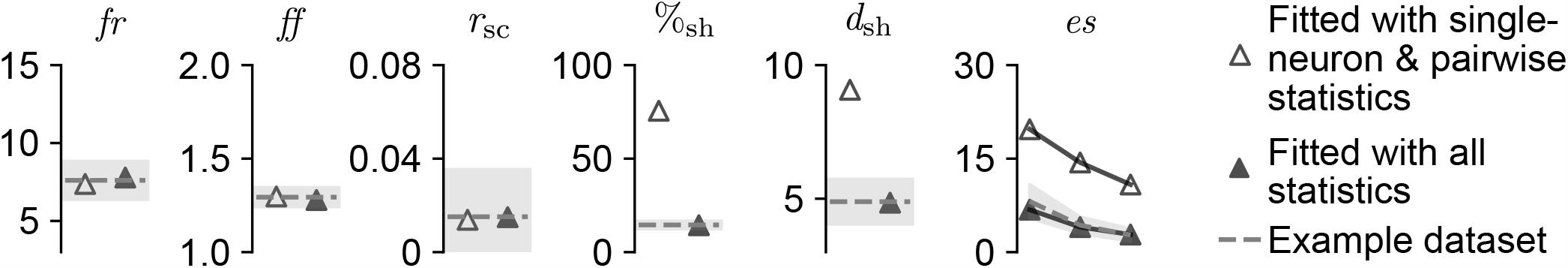
Including population statistics in the cost function improves the quality of fit. In Fig. 6, we found that the SBN is more flexible than the CBN in reproducing activity statistics. When a model is more flexible, it might need to be more strongly constrained during the customization process to accurately reproduce the activity statistics. Here, we ask if the SBN model is well-constrained enough such that simply customizing single-neuron and pairwise statistics will automatically lead to accurately reproducing the population statistics. We applied SNOPS to customize the SBN to an example PFC dataset under two scenarios: (1) only single-neuron and pairwise statistics were included in the cost function (open triangles), and (2) all single-neuron, pairwise, and population statistics were included in the cost function (filled triangles). Although the single-neuron and pairwise statistics were accurately reproduced in both scenarios, leaving the population statistics out of the cost function led to a poor match in population statistics (open triangles are far from the dashed lines). Hence the inclusion of population statistics in the cost function is necessary for constraining the SBN during the customization process. More generally, customizing models even more flexible than the SBN might necessitate the inclusion of additional activity statistics in the cost function.

**Supplementary Figure 8:**
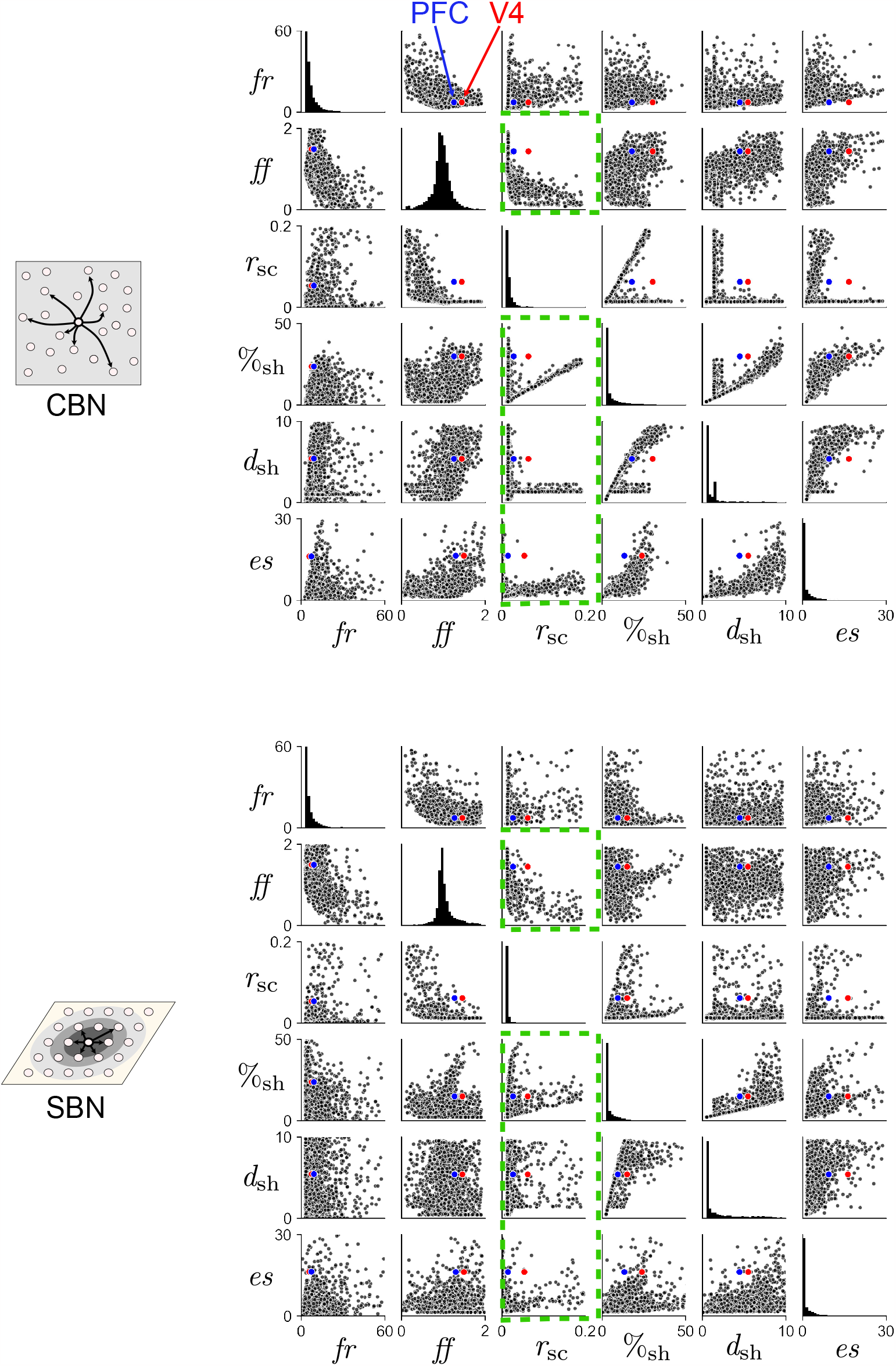
SBN is able to generate a wider combination of statistics than CBN. In Fig. 6, we assessed each model’s flexibility by asking whether it could simultaneously reproduce two or more activity statistics. Here we visualize the range of combinations of activity statistics that each model is able to generate. We randomly drew 5, 000 parameter sets from the search range of each model (see *Spiking network models*). For each parameter set, we simulated activity and computed its activity statistics. Each dot represents a pair of activity statistics for one randomly sampled parameter set. For illustrative purposes, we only show the parameter sets whose activity statistics fall within the following ranges: *fr* : [0, 60], *ff* : [0, 2], *r*_sc_ : [0, 0.2], %_sh_ : [0, 50], *d*_sh_ : [0, 10], *es* : [0, 30]. The red and blue dots represent the example V4 and PFC activity statistics in Fig. 5. The vertical axes of the histograms (subpanels along the diagonal) represent the proportion of dots rather than the vertical axes indicated. The dots visually occupy a greater area for the SBN than CBN, indicating that the SBN is capable of generating more diverse pairs of statistics than the CBN. This is particularly prominent when comparing the CBN and SBN for the subpanels highlighted by the dashed green boxes. In these subpanels, the V4 statistic pairs (red dots) lie outside the range of the statistic pairs generated by the CBN (black dots, upper plot) but inside the range of those generated by the SBN (black dots, lower plot). This demonstrates that the CBN is incapable of simultaneously reproducing *r*_sc_ and any one of the following statistics – *ff*, %_sh_, *d*_sh_, or *es* – of the V4 dataset. This is consistent with the activity trade-off costs identified in Fig. 6c. The visualizations here provide further evidence that the SBN is more flexible than CBN in producing multiple activity statistics simultaneously.

## Notes

### Competing Interest Statement

The authors have declared no competing interest.

